# fMRI signals of pattern separation in the neocortex and hippocampus to non-meaningful objects and their spatial location

**DOI:** 10.1101/2024.10.09.617494

**Authors:** Zsuzsanna Nemecz, István Homolya, Alex Ilyés, Hunor Kis, György Mező, Virág Anna Varga, Markus Werkle-Bergner, Attila Keresztes

## Abstract

Computational theories of memory posit that the dentate gyrus and CA3 (CA3DG) hippocampal subfields reduce mnemonic interference via a process called pattern separation. While the CA3DG is viewed as a domain-general pattern separator, the parahippocampal and perirhinal cortices may play a role in content-specific (e.g., spatial or object-related) interference reduction. Recent work highlighted the role of frontal and parietal control areas in allocating resources during mnemonic discrimination, but the interactions between the medial temporal lobe and frontoparietal regions have been rarely studied. Moreover, mnemonic discrimination tasks designed for humans almost exclusively use everyday items as stimuli, confounding retrieval processes with pattern separation. To address these challenges we acquired high-resolution structural images of the medial temporal lobe, and full-brain high-resolution functional MRI data of 39 participants while they studied non-meaningful fractals with varying degrees of interference in either their spatial or object features. We found that the parahippocampal cortex contributes to interference reduction in the spatial domain, while the perirhinal cortex contributes to interference reduction in the object domain. The dorsolateral frontal and parietal regions were recruited during the encoding of interfering stimuli in both object and location domains, and displayed strengthened within- and cross-network connectivity in response to interference. Contrary to our expectations, we did not find significantly increased activation in the CA3DG to similar trials relative to repeats, indicating a lack of sensitivity to small differences in interfering stimuli. Altogether, these results are in line with content-specific interference reduction in the medial temporal lobe, possibly orchestrated by frontoparietal regions, but challenge the view of the CA3DG as the universal pattern separator of the human brain.

## Introduction

Decades of animal and human research have implicated the dentate gyrus and CA3 subfields of the hippocampus as the primary mnemonic pattern separator of the mammalian brain (Amer & Davachi, 2023; Leal & Yassa, 2018; Leutgeb et al., 2007; Rolls, 2013; Yassa & Stark, 2011). Neuroanatomical accounts propose that divergent excitatory projections from the entorhinal cortex to a large number of sparsely firing neurons expand the available representational space in the dentate gyrus (Cayco-Gajic & Silver, 2019). Consistent with this idea, rodent research has shown that even small changes in the environment induce decorrelated activity patterns in the dentate gyrus (GoodSmith et al., 2019; Knierim & Neunuebel, 2016; Leutgeb et al., 2007; Neunuebel & Knierim, 2014). Moreover, selectively disrupting dentate gyrus function in rats (Gilbert et al., 2001; Hunsaker et al., 2008; McHugh et al., 2007) leads to impaired performance on tests assumed to require spatial pattern separation.

In humans, mnemonic discrimination tasks and functional magnetic resonance (fMRI) imaging provided evidence that levels of lure-related univariate activity in the hippocampus and CA3DG are different from repeats and comparable to new trials (Bakker et al., 2008; Kirwan & Stark, 2007). The parametric manipulation of lure similarity relative to repeats has led to the description of non-linear increases in activity in the hippocampus (Motley & Kirwan, 2012; Stark et al., 2013; Yassa et al., 2011). More recently, studies using multivariate fMRI analysis corroborated the notion that the hippocampus reduces the representational overlap between interfering inputs (Berron et al., 2016; Chanales et al., 2017; Favila et al., 2016; Kyle et al., 2015; Wanjia et al., 2021).

An important and often overlooked confound is that tests of mnemonic discrimination rely on everyday objects or a varying arrangement of these in spatial scenes. Images of everyday objects could induce retrieval of already existing memory representations and confound pattern separation with pattern completion or retrieval processes (Liu et al., 2016). Relatedly, this method cannot differentiate between the orthogonalization of incoming inputs (i.e., true pattern separation) and disparate voxel activity patterns due to partially overlapping, but pre-existing engrams (Quiroga, 2020). Using never-before-seen, non-meaningful stimuli and investigating pattern separation signals at the time of encoding (Liu et al., 2016), could mitigate these concerns by limiting the activation of already available memories and prior associations.

Recent work also raises the question how extra-hippocampal areas contribute to performance in mnemonic discrimination tasks (Amer & Davachi, 2023; Klippenstein et al., 2020; Nash et al., 2021). According to the recent cortical-hippocampal pattern separation framework, frontoparietal regions could play a key role in directing attention to unique features (Amer & Davachi, 2023), and amplifying the separation of memory representations in the medial temporal lobe (MTL). In turn, cortical MTL areas may be engaged in domain-selective interference reduction. Specifically, the perirhinal cortex (PRC) is implicated in processing and discriminating objects (Barense et al., 2010; Berron et al., 2018; Burke et al., 2014; Bussey et al., 2002; Eacott & Gaffan, 2005; Reagh & Yassa, 2014), while the parahippocampal cortex (PHC) is implicated in processing spatial relations and resolving spatial-contextual interference (Berron et al., 2018; Eacott & Gaffan, 2005; Reagh & Yassa, 2014; Staresina et al., 2011). However, the degree of domain-selectivity is debated (Burke et al., 2018; Nilssen et al., 2019), and some studies employing visual and mnemonic discrimination designs did not find such domain specificity (Azab et al., 2014; Lawrence et al., 2020).

The goal of the current study was to assess domain-specific interference reduction in the PHC, PRC, and hippocampus using complex unfamiliar fractals as stimuli in order to reduce the confounding effects of semantic knowledge and retrieval processes. We hypothesized that the PRC will be involved in object encoding, the PHC will be involved in spatial encoding, and the CA3DG will be engaged in interference reduction irrespective of the domain. Further, we conducted a whole-brain analysis followed up by seed-based connectivity analysis to investigate which additional areas are involved in interference reduction of similar stimuli. Although this analysis was exploratory, we expected—based on the cortical-hippocampal pattern separation framework (Amer & Davachi, 2023)—that dorsolateral and parietal regions would be activated by both object and spatial interference, and their connectivity would be modulated by interfering stimuli.

## Methods

### Participants

Power analysis based on pilot behavioral data and conducted with the software GPower indicated a sample size of 40 participants needed to show the behavioral effects of the similarity manipulations at 95% statistical power. Forty-six healthy volunteers (25 female, 21 male, 20–30 years old) were recruited for the study from the local student community. All participants were screened for psychiatric or physical illness, metal implants and current medications. The study was approved by the Ethical Committee of the Medical Research Council, Hungary (ethics approval number: OGYÉI/28291-2/2020), and was carried out in accordance with the code of ethics for human experiments (Declaration of Helsinki). Participants gave written informed consent, and they received either monetary compensation or course credit for their participation. Data of five participants could not be used for the analysis because of technical issues in the MRI (n=3), unfinished data acquisition (n=1), and missing log files due to experimenter error (n=1). Two participants were excluded due to incidental neurological findings. The final sample included 39 participants (19 female, 20 male, 20–30 years old with mean age of 22.61 years and standard deviation of 2.36 years).

### Stimuli

Fractal images (in size 300 * 300 pixels) were generated using Python 3.7.3 by applying the Newton-Raphson method to finding the roots of polynomials and transcendental functions. We created 181 fractal types, each defined by a unique function, and 3 versions of each fractal type (Figure 1), defined by the number of iterations applied, resulting in a stimuli set of 543. Eleven fractal types were used as demonstration and in practice trials, one type was used as a baseline trial that occurred 8 times across tasks, 72 types were randomly assigned to the Location Task, and 96 were randomly assigned to the Object Task. Of the 96 fractal types for the object task, 72 were selected for encoding and 24 served as foils during retrieval (see the detailed description below). Fractal images used in the experiment are available here: https://github.com/zs-nemecz/TerKepEsz/tree/MRI_version/stimuli/fractals. Additionally, an art gallery framing was created by combining drawn images of furniture and plants freely available on the internet.

**Figure 1.**
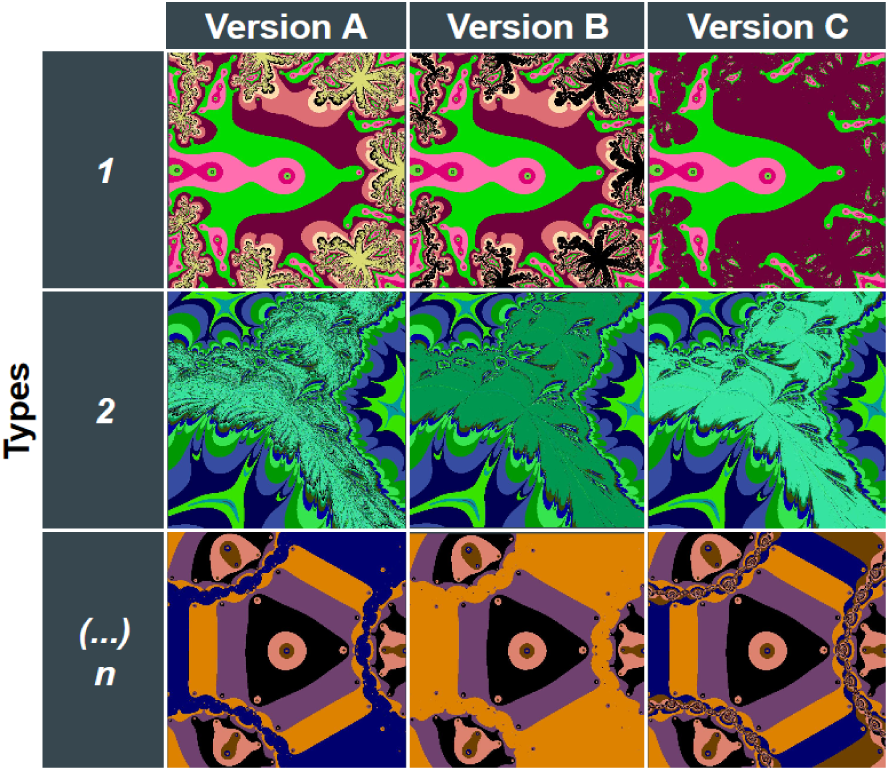
Example fractal stimulus types. Fractals were generated by applying Newton’s method to polynomial and transcendental functions. Fractal types were defined by a unique function and a random color assignment. Three versions of each fractal type – defined by the number of iterations applied – were selected into the stimuli set.

### Experimental design

We used two mnemonic discrimination tasks (the Object Task and the Location Task) with either item or spatial similarity manipulation to test content-dependent interference reduction in the medial temporal lobe. The tasks were created in PsychoPy 2021.1.4 (Peirce et al., 2019). Each task consisted of an encoding and a recognition phase, divided into 4 scanner runs per task (2 encoding runs and 2 recognition runs in each task). During both tasks, a two-dimensional representation of the furnished interior of an art gallery was shown to participants, which contained an empty inner rectangle (1000 by 800 pixels). The location of fractals in both tasks was restricted to the inner rectangle. We used such a framing to enhance the cover story of the task instructions–that participants are sorting and arranging artwork for an exhibition–and to help contextualize spatial locations in the Location Task.

Each trial consisted of a fractal appearing in the inner rectangle of the screen. Trial duration was 3s during the encoding phase, and 4s during the recognition phase. All trials were preceded by a fixation cross, which was presented at the center of the consecutive fractal’s location for a jittered interval between 0.5 and 8s.

After each encoding run and preceding the recognition run, participants had to solve a simple mathematical problem to reduce recency effects, which was followed by a short break. The delay between the last encoding trial and first recognition trial was around 2 minutes on average. Participants were aware of the memory manipulation already during encoding, and they were instructed before the recognition phase to pay attention to small changes of fractals in the Object Task, and small changes of location in the Location Task.

### Object task

The Object task structure is depicted in Figure 2. An overview of stimuli assignment in the Object task is shown in Supplementary Figure 5.

**Figure 2.**
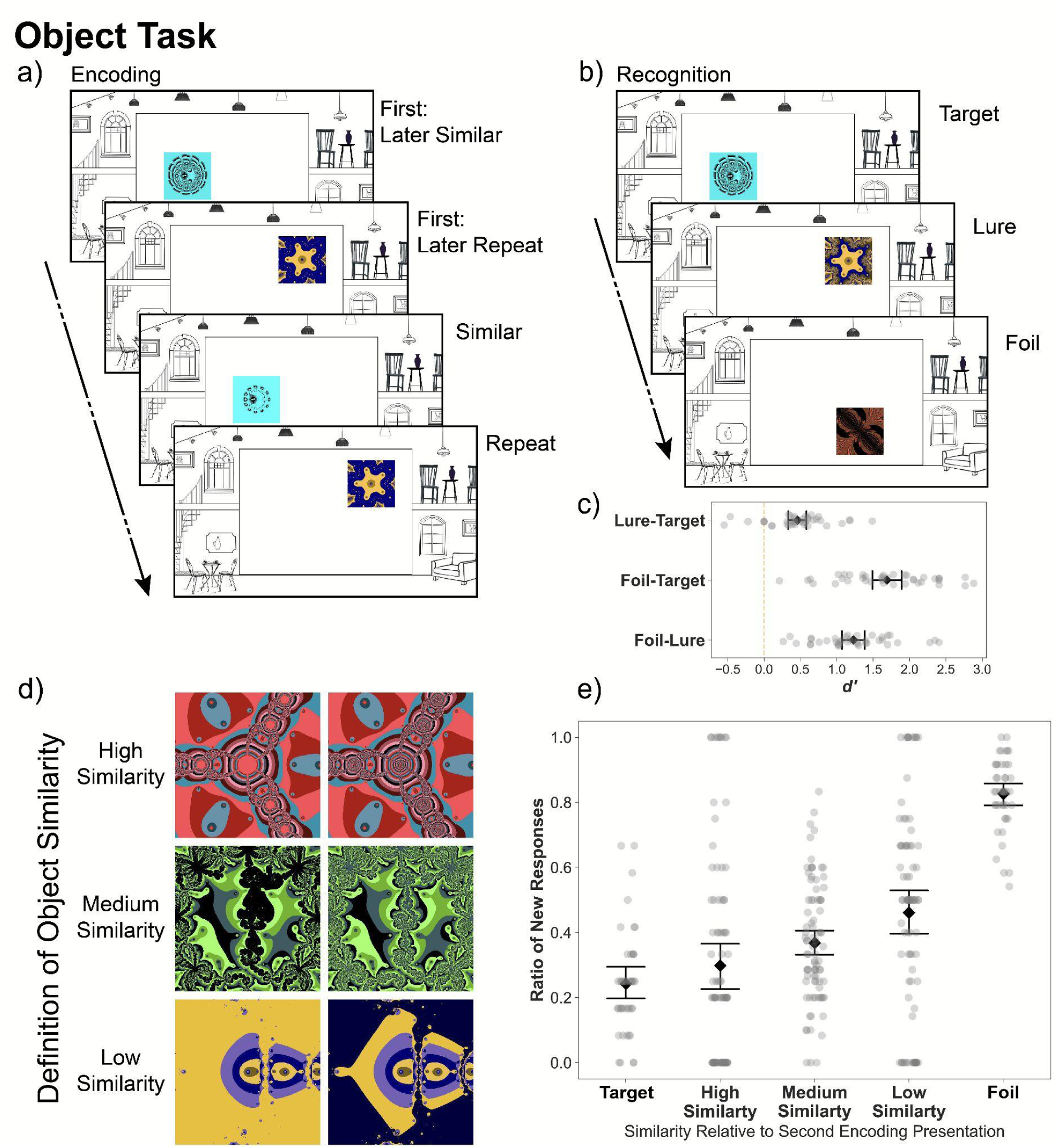
Object Task. Design, procedure, and recognition performance. a) The structure of the Object Task. During encoding, participants saw fractal images appear in an art gallery context (constant through the experiment) and were instructed to make aesthetic judgments about the images. Each trial lasted 3s, with jittered inter-trial intervals (0.5-8s). b) During recognition, participants had to judge whether they had seen exactly this fractal in the previous encoding phase. Trial duration was 4s with jittered inter-trial intervals (0.5-8s). c) Discriminability of each condition in the Object recognition phase. d) Example pairs from each similarity bin used for the behavioral analysis. An independent sample of 30 participants rated each fractal pair on a scale of 0-9 (Identical - Very Different). e) The ratio of “new” responses is dependent on the similarity of lure fractals. The x axis depicts the similarity between the first presentation of the fractal type and the version presented at recognition. The y axis depicts the raw ratio of “new” responses for each participant. Error bars represent 95% confidence intervals.

#### Encoding phase

In the Object encoding phase (Figure 2a) participants made binary aesthetic judgements about each fractal, answering the following question: “Would you like to display this image in the exhibition of the art gallery?”. This question served two purposes. First, we wanted to make sure participants attend to the features of each fractal. Second, although in principle passive viewing could have elicited our neural signals of interest, by requiring a response from participants, we could make sure that they pay attention throughout the task. For encoding, we selected 72 fractal types that were shown in one of the three possible versions (VER_1st_) at their first appearance at a randomly assigned location (LOC_1st_). We refer to those first presentations as new stimuli at encoding (NEW_ENC_). From those 72 fractal types, 24 were repeated during encoding in the same version at exactly the same location (LOC_1st_). We refer to those second presentations as repeats during encoding (REP_ENC_). The remaining 48 fractal types were also shown a second time during encoding at the same location (LOC_1st_), but in a different version (VER_2nd_). We refer to those second presentations as similar during encoding (SIM_ENC_). We used twice as many similar items as repeats to increase the power of detecting differences between similarity bins, and small increases relative to repeats, assuming that repetition suppression is a robust effect that may be detectable with less trials than the sensitivity to similarity. The lag between first (NEW_ENC_) and second presentations (SIM_ENC_ and REP_ENC_ trials) was jittered between 2 to 6 trials.

#### Recognition phase

In the Object recognition phase (Figure 2b) participants had to make “old”/”new” memory judgements, corresponding to the following question: ‘Have you seen exactly this image before?’. Twentyfour of the recognition trials were targets (same fractal at the same location as the original), 24 were lures (another version of the same fractal type, presented at the same location as the original), and 24 were foils (new fractal at a randomly assigned location). For the recognition phase we selected 48 fractal types from the first presentations during encoding (i.e., NEW_ENC_). Half of the fractal types, i.e., 24, became target items during recognition (TARGET_REC_). The other half, i.e., 24, was used as lure items during recognition (LURE_REC_). During recognition, TARGET_REC_ items were always shown in the exact same version they appeared during the first presentation at encoding (i.e., VER_1st_). The 24 TARGET_REC_ items were selected in a way that 12 have been repeats during their second presentation at encoding (i.e., REP_ENC_), while the remaining 12 have been shown in a similar version during their second presentation at encoding (i.e., SIM_ENC_). The LURE_REC_ items were always shown in the fractal version of the three possible ones that was not yet used in the task, so far (i.e., VER_3rd_). Similar to the target items, also the 24 LURE_REC_ items were selected in a way that 12 have been shown as exact repeats (i.e., REP_ENC_) and 12 as similar versions (i.e., SIM_ENC_) during their second presentation at encoding. The 24 target and 24 lure items were intermixed with 24 foil items (FOIL_REC_). The FOIL_REC_ items were selected from novel fractal types that were not presented during encoding. The version of FOIL_REC_ fractals was radomly drawn from the three possible versions (i.e., VER_foil_).

#### Object similarity manipulation

Similar (SIM_ENC_) and lure (LURE_REC_) fractals in the Object Task were different versions of the same fractal type as the original fractal presented during encoding (VER_1st_, Figure 1), i.e., the fractal was created using the same function but a different number of iterations. The similarity between fractal versions was rated post-hoc by an independent sample of participants (n=30) on a scale from 0 (“Same”) to 9 (“Very Different”). Details of the rating experiment and the distribution of fractal similarity values are included in the Supplementary Materials. Figure 2d shows examples of fractal pairs with high, medium and low similarity ratings.

### Location task

The Location task structure is depicted in Figure 3. An overview of stimuli assignment in the Location task is shown in Supplementary Figure 6.

**Figure 3.**
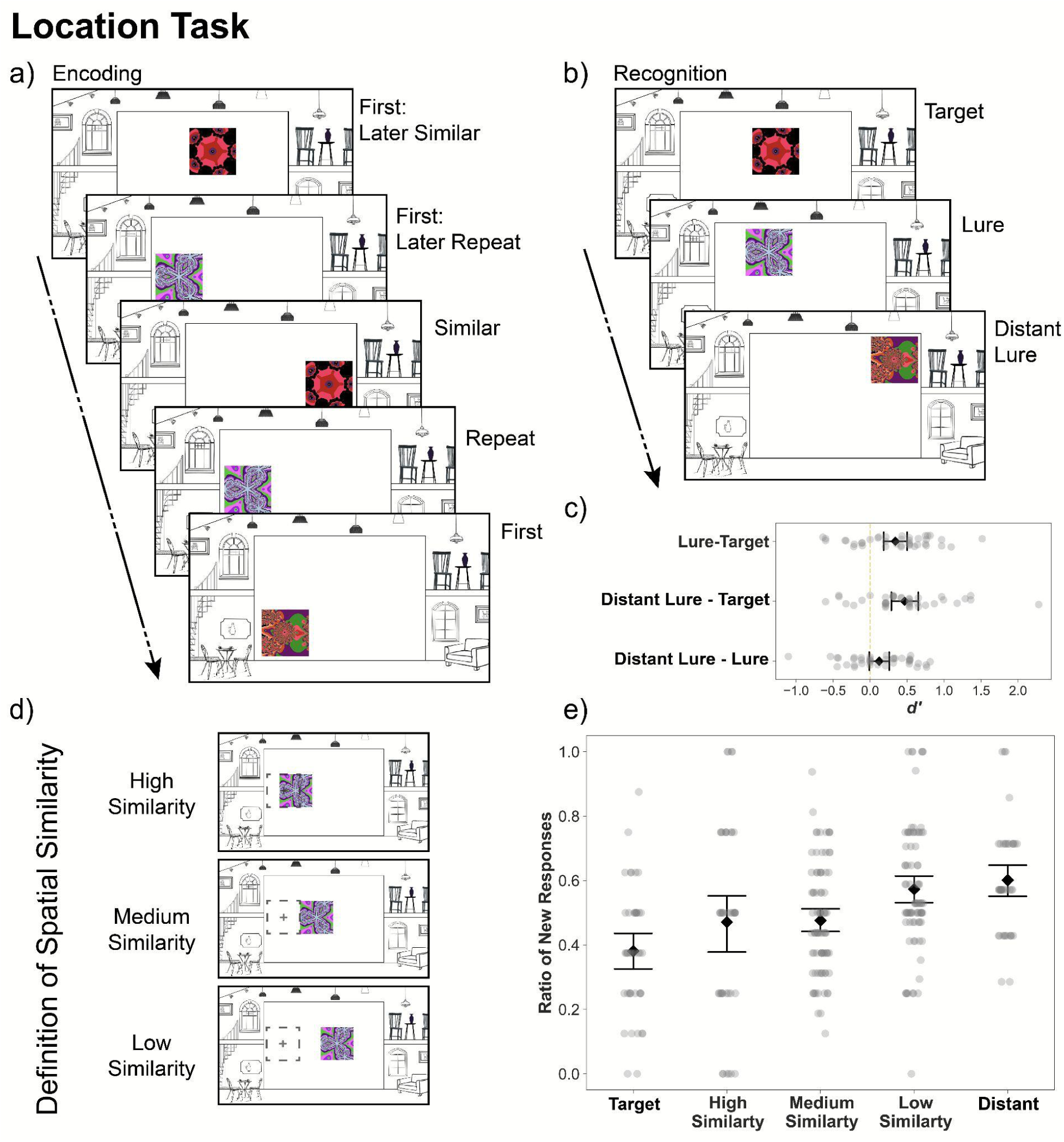
Location Task. Design, procedure, and recognition performance. a) The structure of the Location Task. During location encoding, participants saw fractal images appear in an art gallery context (constant through the experiment) and were instructed to make aesthetic judgments about the position of the images. Each trial lasted 3s, with jittered inter-trial intervals (0.5-8s). (b) During recognition, participants had to judge whether they had seen the shown fractal at the exact current location in the preceding encoding phase. Trial duration was 4s with jittered inter-trial intervals (0.5-8s). c) Discriminability of each condition in the Location recognition phase. d) Example pairs from each similarity bin used for the behavioral analysis. Very close pairs were presented 1-200 pixels away, close pairs were presented 200-400 pixels away, far pairs were presented 400-600 pixels away from the second presentation. Very far pairs are more than 600 pixels away from the second presentation. e) The ratio of “new” responses is modulated by distance. The x axis depicts the spatial similarity between the second presentation of the trial and the recognition trial. The y axis depicts the raw ratio of “new” responses for each participant. Error bars represent 95% confidence intervals.

#### Encoding phase

The encoding phase of the Location Task (Figure 3a) was analogous to the encoding phase of the Object Task, but here similar trials were defined by their Euclidean distance to the first presentation of the fractal, not their within-item features. Participants made binary aesthetic judgments about the location of each fractal image. Specifically, participants had to decide whether they would like to display the presented image at the current location in the art gallery, which encouraged participants to pay attention to the location of each fractal. For encoding, we selected 72 fractal types that were shown in one of the three possible versions (VER_1st_) at their first appearance at a randomly assigned location (LOC_1st_). We refer to those first presentations as new stimuli at encoding (NEW_ENC_). From those 72 trials, 23 were repeated during encoding in the same version (VER_1st_) at exactly the same location (LOC_1st_). We refer to those second presentations as repeats during encoding (REP_ENC_). The remaining 49 fractals were also shown a second time during encoding in the same fractal version (VER_1st_), but at an alternative location (LOC_2nd_). We refer to those second presentations as similar during encoding (SIM_ENC_). Locations were unique to each NEW_ENC_ and SIM_ENC_ trial, thus there were altogether 120 locations used in the Location encoding phase. Trial order was pseudo-randomized, and the lag between first (NEW_ENC_ trials) and second presentations (SIM_ENC_ and REP_ENC_trials) was jittered between 1 to 6 trials.

#### Recognition phase

During the Location recognition phase (Figure 3b), fractals were presented again in the inner rectangle of the screen, and participants had to make “old”/”new” memory judgements, corresponding to the following question: ‘Have you seen this image exactly at this location?’. Participants were made aware of the fact that only fractals shown in the previous encoding phase were presented during recognition. For the recognition phase we used all 72 first presentation trials from the encoding phase (i.e., NEW_ENC_). One third of these trials, i.e., 24, was used as target trials (TARGET_REC_), one third served as the basis for lure trials (LURE_REC_), and one third for distant lure trials (DISTANT_REC_). TARGET_REC_ trials were always shown in the exact same fractal version and exact same location as they appeared during the first presentation at encoding (i.e., VER_1st_ LOC_1st_). The 24 TARGET_REC_ trials were selected in a way that 8 have been repeats during their second presentation at encoding (i.e., REP_ENC_), while the remaining 16 have been shown at a similar location during their second presentation at encoding (i.e., SIM_ENC_). The LURE_REC_ trials were never presented in an exact fashion during encoding, but the same fractal (VER_1st_) was presented at a different location during encoding (i.e., LOC_3rd_). Similar to the target items, also the 24 lure items were selected in a way that 8 have been shown as repeats (i.e., REP_ENC_) and 16 as similar (i.e., SIM_ENC_) during their second presentation at encoding. The distant lure trials (DISTANT_REC_) were VER_1st_ fractals presented in a location at least 450 pixels away (LOC_DIST_) from the first presentation location (LOC_1st_). TARGET_REC_, LURE_REC_, and DISTANT_REC_ trials were intermixed and their order was pseudorandomized.

#### Spatial similarity manipulation

During the encoding phase of the Location Task, similar trials (SIM_ENC_) consisted of repeated (VER_1st_) fractals presented 200 to 350 pixels away from their original location (LOC_2nd_). The exact distance was randomly sampled from a uniform distribution.

For the recognition phase, we originally grouped trials into target (TARGET_REC_), lure (LURE_REC_) and distant lure (DISTANT_REC_) conditions based on their Euclidean distance to the very first presentation of the fractal in the encoding phase (LOC_1st_). Twentyfour of the recognition trials were thus exactly repeated, twentyfour were lures separated by at least 200 and maximum 350 pixels, and 24 were distant lures separated by at least 450 pixels. However, each trial’s spatial similarity can also be calculated based on the Euclidean distance to the second presentation of the fractal during encoding, which also affected participants’ “new” responses (Fig. 3e). Distributions of lure and distant lure distances to the first and second presentation can be found in the supplementary materials (Suppl. Fig. 7).

### Experimental procedure

All participants completed both the Location and the Object Task. The order of the tasks was counterbalanced across participants. Participants were taken out of the scanner between the tasks for a longer break, and they were given the task-specific instructions immediately preceding the task. Before each task, participants were given 6 practice trials with feedback. Encoding and recognition phases were divided into two runs each, and participants completed them in an interleaved fashion, so that a recognition run followed each encoding run. For example, a participant starting with the Object Task had the following procedure: Object Task Instructions and Practice, Object Encoding 1, Object Recognition 1, Self-paced break in the scanner, Object Encoding 2, Object Recognition 2, Long break outside the scanner, Location Task Instructions and Practice, Location Encoding 1, Location Recognition 1, Self-paced break in the scanner, Location Encoding 2, Location Recognition 2.

### Behavioral data analysis

We analyzed the ratio of “new” responses with a one-way repeated-measures ANOVA with the within factor condition (target, lure, foil) in Python 3.9.13 with statsmodels 0.13.2 (Seabold & Perktold, 2010). Follow-up paired sample t-tests were conducted with scipy 1.13.1 (Virtanen et al., 2020), and we controlled for the false discovery rate (FDR) with the Benjamini-Hochberg method (Benjamini & Hochberg, 1995). Figures were created with matplotlib 3.9.2 (Hunter, 2007) and seaborn 0.13.0 (Waskom, 2021).

To further assess the effect of similarity per each trial, we used generalized binomial linear mixed effects modeling implemented in the R package lme4 1.1-31 (Bates et al., 2015). For each task, we built 3 models, predicting the probability of a “new” response per recognition trial. Assumption checks for the mixed effects models are described in the supplementary.

The first model (Model A) included the similarity between the first encoding trial and the recognition trial as a fixed effect predictor, and the participant-specific intercept as a random effects predictor. The second model (Model B) included the similarity between the second encoding trial and the recognition trial as a fixed effect predictor, and the participant-specific intercept as a random effects predictor. The third model (Model C) included the similarity between both the first and the second encoding trial and the recognition trial as fixed effect predictors, and the participant-specific intercept as a random effects predictor. Accordingly, for each task the three models were defined as below:

Model A: RespNew ∼ FirstSimilarity + (1 | Participant),
Model B: RespNew ∼ SecondSimilarity + (1 | Participant),
Model C: RespNew ∼ FirstSimilarity * SecondSimilarity + (1 | Participant),

where *RespNew* is the probability of a “new” response, *FirstSimilarity* is the similarity between the first encoding trial and the recognition trial, and *SecondSimilarity* is the similarity between the second encoding trial and the recognition trial.

For the correlations between behavior and neural measures, we obtained the measures of individual sensitivity to similarity from modified versions of models A and B, which, in addition to the random intercepts, included random slopes. This type of model allows for the relationship between the explanatory variable (here: similarity) and the predicted variable (here: new response) to be different for each individual. Participant specific coefficients (individual slopes) were then extracted from the models.

### MRI data acquisition

A Siemens Magnetom Prisma 3T MRI scanner (Siemens Healthcare, Erlangen, Germany) with the standard Siemens 32-channel head coil was used. Functional measurements were acquired with a conventional gradient-echo-EPI sequence using multiband-acceleration with factor 4 (80 slices, slice gap: 0.375 mm, slice ordering: interleaved, phase encoding direction: anterior to posterior, repetition time (TR) = 1840 ms, echo time (TE) = 30 ms, flip angle (FA) = 72°; field of view (FOV) = 204 mm, spatial resolution 1.5 × 1.5 × 1.5 mm). Fieldmaps for the functional runs were acquired each time the participant was placed in the scanner, resulting in one fieldmap per task. We obtained sagittal T1-weighted images using a 3D MPRAGE sequence (TR = 2300 ms,TE = 3.03 ms, FA = 9°, FOV = 256 mm, spatial resolution 1 × 1 × 1 mm). An additional restricted FOV high-resolution proton-density (PD) weighted image of the medial temporal lobe area was acquired (TR = 6500 ms, TE = 16 ms, FA = 120°; spatial resolution 0.4 × 0.4 ×2 mm, FOV = 206 mm) to aid the delineation of the anatomical regions of interest.

### Subject-level ROI analysis

Only data from the encoding runs are analyzed here. Functional data preprocessing and analyses were done using FSL FEAT (Jenkinson et al., 2012; Woolrich et al., 2001, 2004, 2009). Functional runs were undistorted using fieldmaps and the T1 structural image. Then, each slice was spatially realigned to the middle slice within each volume with FSL MCFLIRT (Jenkinson et al., 2002), resulting in motion parameters that were used as regressors in further analysis for motion correction. As we strived to retain as much individual anatomical specificity as possible, functional runs were not transformed to the T1 or standard space for the region of interest analysis (see section “Registration of Anatomical Regions of Interest” below). In order to enable the analysis of multiple runs per subject in a single higher-level model, we used affine registration with 6 degrees of freedom to transform each run per participant to the same participant-specific functional space. To take advantage of the high spatial resolution functional scans, and considering the small size of hippocampal subfields, we applied minimal smoothing on the data with a 2-mm full-width at half-maximum isotropic Gaussian kernel. Finally, we applied a high-pass filter with a threshold of 90s on functional runs. The threshold of 90s was estimated by FSL Feat based on the trial timing.

For each participant, we performed two general linear model analyses using the data of the encoding phase. Regressors were defined by convolving the canonical hemodynamic response function with a boxcar function representing the onsets and duration of the events within each explanatory variable.

First, we modeled the new, similar, and repeated encoding trials in a 2 (presentation rank: first vs second) - by - 2 (trial type: similar vs. repeat) design with four regressors: first presentation of later repeats (Presentation 1, Type Repeat), first presentation of later similar trials (Presentation 1, Type Similar), repeats (Presentation 2, Type Repeat), and similar trials (Presentation 2, Type Similar). The design is depicted in Figure 4a and the interpretation of expected results is depicted in Figure 4b. We opted for this balanced model to ensure that differences between Types are not confounded by potential baseline differences between the first presentations of later repeat and similar trials. In a 2-by-2 repeated measures ANOVA, such an imbalance would be indicated by a main effect of trial type. We found no indication of a baseline difference between trial types, and thus we collapsed all first presentation trials in the follow-up analysis.

**Figure 4.**
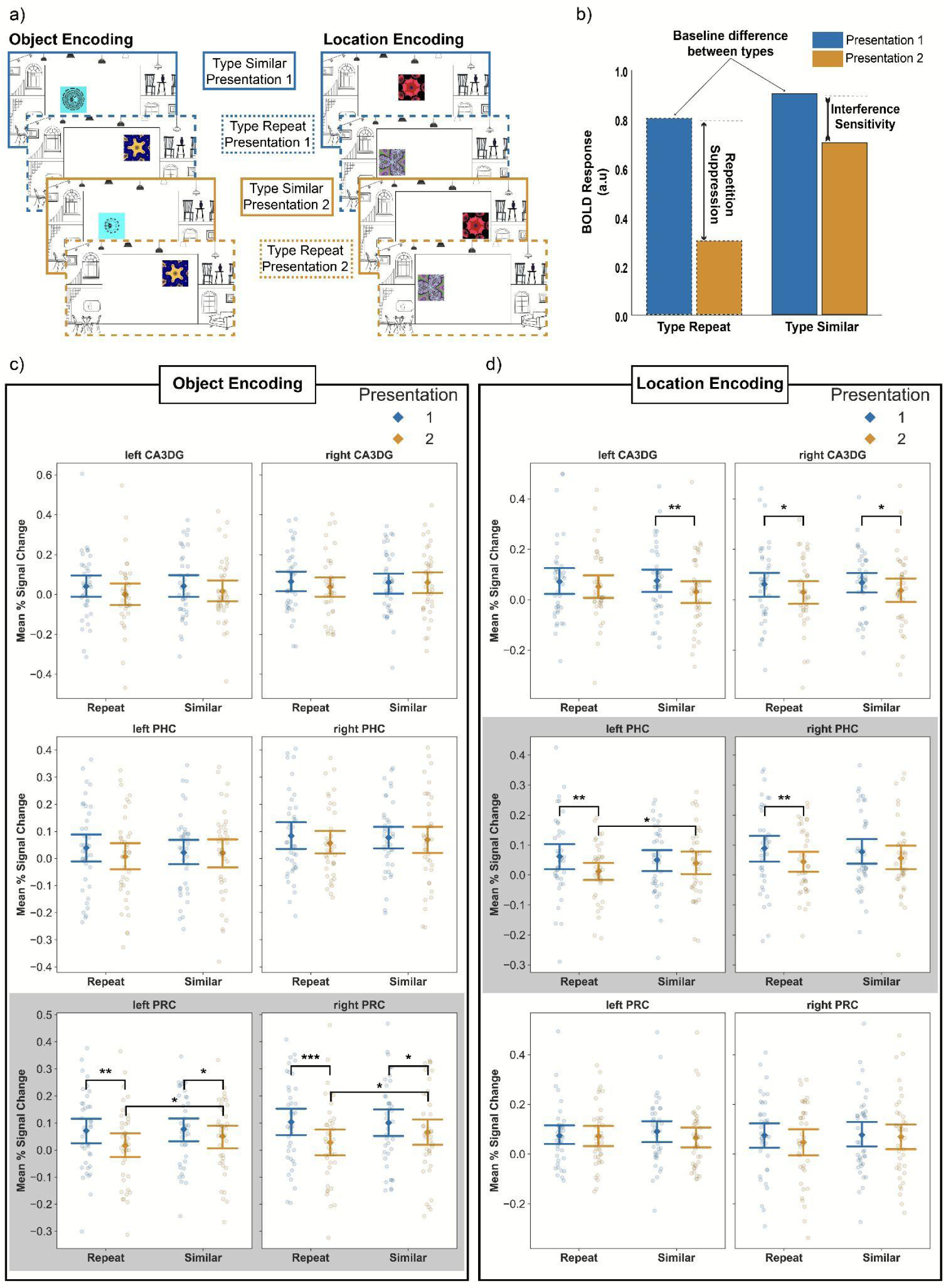
BOLD responses in the encoding phase modeled in a 2-by-2 design. a) First presentation trials were divided into first presentations of later similar trials and later repeats. b) Expected results and their interpretation from the 2-by-2 model. c) Responses of the main medial temporal regions of interest during the object encoding. d) Responses of the main medial temporal regions of interest during location encoding. Neural response patterns in c) and d) suggesting differentiation between similar and repeat trials are shaded with gray. Significant differences (unadjusted) are denoted with a symbol: * p < .05, ** p < .01, *** Abbreviations: CA3DG - CA3 and dentate gyrus, PHC - parahippocampal cortex, PRC - perirhinal cortex. Error bars represent 95% confidence intervals.

**Figure 5.**
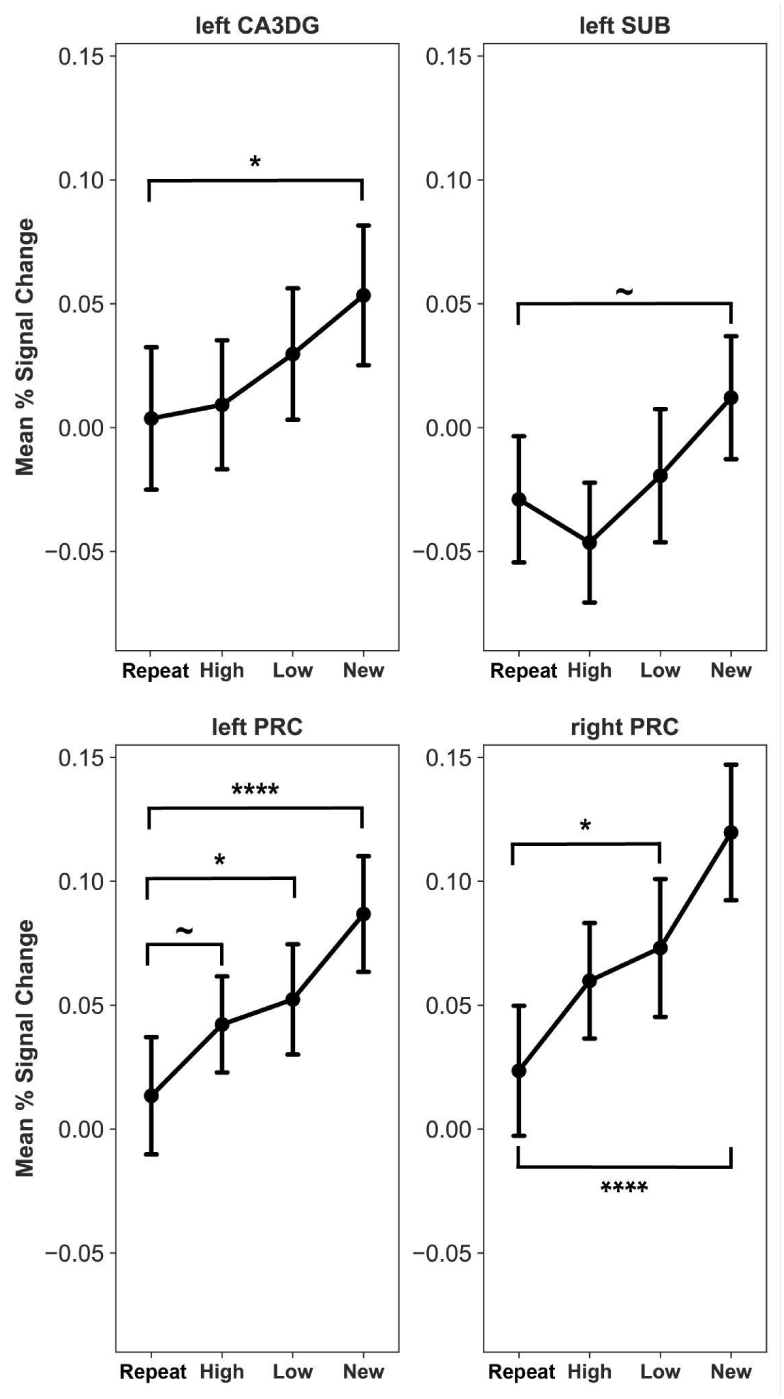
Similarity dependent activations during object encoding. The PRC expresses a significant increase to low similarity object trials bilaterally. Significant differences relative to repeats (unadjusted) are denoted with a symbol: ∼ p < .06, * p < .05, ** p < .01, *** p < .001, **** p < .0001. Responses from all areas that showed a significant effect in the previous 2-by-2 model were included in the figure. Abbreviations: CA3DG - cornu Ammonis 3 and dentate gyrus, SUB - subiculum,, PRC - perirhinal cortex. Error bars represent the standard error of mean.

In the second model, we collapsed all first presentation trials, and divided similar trials into high-similarity and low-similarity trials. This resulted in four explanatory variables of interest: repeat, high-similarity, low-similarity, and new (i.e., first presentation) trials.

A single baseline trial was repeated eight times across tasks. We included the first presentation of this trial among the first presentation of later repeat trials (and in the second model, among the merged first presentation trials), the second presentation among repeats, and the third to the eighth presentations were included in a separate explanatory variable in both models. We deemed the number of these multiple repetition trials too low to include them in further analysis, but exploratory results can be found in Supplementary Figure 2.

In both GLM analyses, we included the six head motion parameters estimated in the realignment procedure and trial-wise reaction times (Mumford et al., 2024) as nuisance regressors. The resulting regressors were fitted to the observed functional time series of the encoding phase. Mean percent signal change values from all regions of interest were extracted from the individual participant space with FSL featquery and were analyzed with Python libraries scipy (Virtanen et al., 2020) and statsmodels (Seabold & Perktold, 2010), and R package lme4 (Bates et al., 2015). All reported *p* values, unless otherwise stated, are FDR adjusted per group of analysis (Benjamini & Hochberg, 1995), to account for multiple comparisons.

### Delineating anatomical ROIs in the MTL

Anatomical regions of interest (ROIs) were delineated on the high-resolution PD images with the software Automatic Segmentation of Hippocampal Subfields (ASHS, Yushkevich et al., 2015). We used the UPENN PMC Atlas (Yushkevich et al., 2015) for the segmentation of the perirhinal and parahippocampal cortices, and a custom atlas (Bender et al., 2018) for the delineation of subfields within the hippocampal body (cornu Ammonis 1-2: CA1, subiculum: SUB, cornu Ammonis 3 and dentate gyrus: CA3DG). Due to the less uniform structure of the hippocampal head and tail, no histologically validated segmentation protocols have yet been established for these regions (Bender et al., 2018; Wisse et al., 2017). Therefore, only the hippocampal body was segmented in this study. As in numerous previous high-resolution fMRI studies (Nash et al., 2021; Reagh & Yassa, 2014), the subfields CA3 and DG were combined into a single ROI (CA3DG). The quality of the segmentations was inspected by an experienced rater (A.K.), and errors were corrected following the recent guideline by Canada et al., 2023. Segmented regions were binarized and used as masks for extracting mean percent signal change values in the univariate and ROI-to-ROI connectivity analysis. An example segmentation is shown in Supplementary Figure 8.

### Registration of functional and high-resolution anatomical images

We defined each participant’s individual functional space as the mean volume of the first Object Encoding run. All other functional runs were registered to this volume with rigid body registration as implemented in FSL FLIRT (Jenkinson et al., 2002; Jenkinson & Smith, 2001) during the analysis of fMRI data. Anatomical masks were transformed into functional space with Advanced Normalization Tools (ANTs, Avants et al., 2009, 2011) using a three-step registration procedure. All input images were N4 bias-field corrected before registration. First, we registered the standard T1 anatomical image of each participant to the corresponding high-resolution partial field-of-view PD image with affine transformations, and obtained the transformation matrix *T1-to-PD*. Second, we registered the mean skull-stripped functional volume to the skull-stripped T1 anatomical image first using rigid transformation and then symmetric image normalization (SyN, Avants et al., 2008), and obtained the transformation matrix *meanFunc-to-T1* and warp image *meanFunc-to-T1*. For the meanFunc to T1 registration, we used a generous medial temporal lobe mask to optimize the registration outcome in this region. Exact parameters of the ANTs registration calls as well as registration examples can be found in the supplementary materials. Third, we transformed medial temporal ROIs derived from the PD space to functional space by applying the inverse of the *meanFunc-to-T1* matrix, the inverse of the *meanFunc-to-T1* warp, and finally the inverse of the *T1-to-PD* matrix.

### Whole-brain group-level fMRI analysis

We generated a group specific anatomical template with ANTs by registering individual T1 scans to a template space and then averaging them to form a whole-brain image representative of our sample. The average of all original T1 input images was used as the starting point for the iterative registration process. N4 bias field correction was applied on all input images before registration. One linear registration of input files was followed by four nonlinear SyN iterations. Each SyN registration was performed over four increasingly fine-grained resolutions (100 × 100 × 70 × 20 iterations). The similarity metric was cross-correlation. Template update steps were set to 0.2 mm.

The fMRI data processing steps for the whole-brain analysis were identical to those of the subject-level region-of-interest analysis described above, with the exception of registering functional runs to the individual T1 scans with BBR registration implemented in FSL (Greve & Fischl, 2009). Individual T1 scans were normalized to the group template with nonlinear registration to facilitate group-level analysis.

We modeled first, similar, and repeated encoding trials in a 2-by-2 design with four regressors as above (Figure 4a), separating the first presentation of later similar and of later repeat trials into two regressors. Following previous work (Nash et al., 2021; Reagh & Yassa, 2014), we chose the contrast *similar - first presentation of later similar trials* as our comparison of interest to identify areas that show high activity to similar trials, disregarding the visual features they share with first presentations and tapping on mnemonic processes. Group-level contrast parameter estimate (COPE) images were transformed into MNI152 standard space, after coregistering the group T1 template image to the 1mm MNI152 template brain available in FLS. We obtained anatomical labels reported in the whole-brain group level results from the Harvard-Oxford Cortical Atlas (Desikan et al., 2006; Frazier et al., 2005; Goldstein et al., 2007; Makris et al., 2006).

### Generalized psychophysiological interaction analysis

Psychophysiological interaction (PPI) analysis examines task-specific modulations of the functional connectivity of an area (Di et al., 2021; Friston et al., 1997) by modeling the interaction between the timeseries of the area (physiological variable) and a task condition (psychological variable). In a generalized psychophysiological interaction analysis, in contrast to the standard PPI approach, each task condition and its interaction with the physiological variable is modeled, thus covering the entire experimental space, leading to better model fit (McLaren et al., 2012).

We selected three areas per hemisphere that emerged consistently across the two tasks during similar lure trials in the whole-brain analysis (the angular gyrus, the supramarginal gyrus, and the middle frontal gyrus) to serve as seeds in the gPPI analysis. Cortical segmentation based on the Destrieux atlas (Destrieux et al., 2010; Fischl, 2004) were acquired for each participant using Freesurfer (Dale et al., 1999; Desikan et al., 2006; Fischl et al., 1999, 2002). Freesurfer segmentations were transformed into functional space, and ROI labels were extracted and binarized. Using the ROI masks, we derived the mean timeseries data from each functional run, and used these as physiological variables. The gPPI models included a regressor for each task condition and the interaction between the seed activity and task condition.

### ROI-to-ROI connectivity analysis

We extracted the mean preprocessed timeseries data from each functional run using our medial temporal (left and right CA3DG, PHC, PRC) and frontal (left and right middle frontal gyrus) and parietal (left and right angular gyrus, supramarginal gyrus) masks. Motion parameters were regressed out of all ROI timeseries data. Next, we correlated the timeseries data from each medial temporal region with all frontal and parietal regions, and created one correlation matrix per functional run and subject. We applied Fisher’s z transformation on the correlation coefficients, and then averaged matrices across runs of the same task, thus acquiring one correlation matrix per subject and task. We tested whether there is a significant connectivity between pairs of regions with one-sample t-tests. Connectivity differences between the Location and Object Encoding were tested with paired sample t-tests. The relationship between connectivity and behavior was assessed with Spearman correlations, as we had no predictions on the linearity of the associations. Pearson correlations yielded the same pattern of results, but indicated stronger relationships between variables.

## Results

### Behavioral results - Recognition phase

#### Ratio of “new” responses per condition

A one-way repeated-measures ANOVA revealed differences in the ratio of “new” responses during the Recognition Phase as a function of condition (target, lure, foil/distant lure) in both tasks (Object: *M_targe_*_t_ = 0.28, *SD* = 0.13, *M_lure_* = 0.43, *SD* = 0.14, *M_foil_* = 0.83, *SD* = 0.1, *F*(2, 76) = 264.99, *p* < .001, *η^2^_generalized_* = 0.78, Location: *M_targe_*_t_ = 0.41, *SD* = 0.13, *M_lure_* = 0.54, *SD* = 0.14, *M_distant_* = 0.58, *SD* = 0.11, *F*(2, 76) = 19.63, *p* < .001, *η^2^_generalized_* = 0.25). We then compared the ratio of “new” responses between conditions directly with paired sample t-tests. As a reminder, all reported *p* values of the post-hoc comparisons are FDR adjusted (Benjamini & Hochberg, 1995). In the Object Task, post-hoc paired sample t-tests showed a significant difference between each combination of conditions (target - lure: *t*(38) = −7.09, *p* < .001, target - foil: *t*(38) = −19.47, *p* < .001, lure - foil: *t*(38) = −16.63, *p* < .001). In the Location Task, post-hoc paired sample t-tests showed a significant difference between target and lure trials (*t*(38) = −4.39, *p* < .001), and target and distant lure trials (*t*(38) = −5.59, *p* < .001), but not between lure and distant lure trials (*t*(38) = −1.69, *p* = .099). Importantly, the significant differences in the ratios of “new” responses between targets and lures, and targets and foils/distant lures in both tasks demonstrated that participants discriminated between previously seen and changed trials.

#### Discriminability (d’) of conditions

Following previous work with a two-response (old/new) version of the mnemonic discrimination task (Loiotile & Courtney, 2015; Stark et al., 2015), we replicated the above results using a signal detection measure of discriminability (*d’*). A *d’* between each pair of conditions revealed that the discriminability of all combinations of conditions was significantly higher than 0 in the Object Task (lure-target: *M_lure-target_* = 0.46, *SD* = 0.4, *t*(39) = 7.00, *p* < .001, foil-lure: *M_foil-lure_* = 1.23, *SD* = 0.5, *t*(39) = 15.12, *p* < .001, foil - target: *M_foil-target_* = 1.7, *SD* = 0.63, *t*(39) = 16.47, *p* < .001). In the Location task it was significantly higher than 0 between lures and targets (*M_lure-target_* = 0.34, *SD* = 0.49, *t*(39) = 4.29, *p* < .001), and distant lures and targets (*M_distant-target_*= 0.47, *SD* = 0.55, *t*(39) = 5.2, *p* < .001), but not distant lures and lures (*M_distant-lure_*= 0.12, *SD* = 0.42, *t*(39) = 1.81, *p* = .078).

#### The effect of trial similarity on the probability of a “new” response

In the above analyses, we assigned the types target, lure and foil/distant lure based on the similarity between the recognition trial and the first encoding trial in which the fractal was presented. However, as each fractal was presented twice during encoding (either in an exactly repeated or similar version, see Figure 2a and Figure 3a), we hypothesized that the similarity between the recognition trial and the *second* presentation of the trial will also modulate participants’ performance. In addition, the ANOVAs presented above disregard trial level similarity information. Therefore, in a follow-up analysis, we calculated the item and spatial similarity between each recognition trial and its first and second encoding presentation, and we assessed the relationship between each similarity measure and the probability of a “new” response using binomial generalized linear mixed effects modeling with random intercepts (see Methods).

The probability of a “new” response in the Object Task was significantly associated with the first presentation item similarity (*p* < .001, Supplementary Table 1 - Model A) and the second presentation item similarity (*p* < .001, Figure 2c, Supplementary Table 1 - Model B). Adding both similarity measures to the model (Supplementary Table 1 - Model C) improved the fit significantly, and resulted in a slightly lower Akaike Information Criterion (AIC, Supplementary Table 2). In this model, only the effect of the similarity between the second presentation and the recognition trial remained significant (*p* < .001).

The probability of a “new” response in the Location Task was significantly associated with the first (*p* < .001, Supplementary Table 3 - Model A) and second (*p* < .001, Supplementary Table 3 - Model B) presentation’s spatial similarity (Figure 3c). Adding both similarity measures to the model (Supplementary Table 3 - Model C) improved the fit, and resulted in a slightly lower AIC value (Supplementary Table 4). In this model, both the first presentation similarity (*p* < .001), second presentation similarity (*p* = .006), as well as their interaction (*p* < .001), were significantly associated with the probability of a “new” response.

### Object encoding: Univariate ROI results

#### Presentation rank by trial type model

In the Object task, a region-wise 2-by-2 repeated measure ANOVA of presentation rank (first and second) and trial type (similar and repeat) revealed a main effect of presentation rank in the following regions: left SUB (*F*(1, 38) = 7.91, *p* = .023, *η^2^_partial_* = 0.17), left PRC (*F*(1, 38) = 16.07, *p* = .002, *η^2^_partial_* = 0.3), and right PRC (*F*(1, 38) = 25.35, *p* < .001, *η^2^_partial_* = 0.4). The effect in the left CA3DG was not significant after adjusting for the FDR (left CA3DG (*F*(1, 38) = 5.79, *p* = .053, *p_unadjusted_* = .021, *η^2^_partial_* = 0.13). The main effect of presentation rank was due to the lower activity level at Presentation 2 compared to Presentation 1 in all regions, i.e. a repetition suppression effect. We did not find a main effect of trial type (all *p*s > .07), or any significant difference between first presentation of similar trials and first presentation of repeat trials in any area (all *p*s > .2). Notably, we did not find any significant effect in either the left or right PHC (all *p*s > .1), indicating that the experimental manipulations in the Object Task left the PHC largely unaffected. Based on our a priori hypothesis that the CA3DG and PRC are involved in object pattern separation, which should result in higher activity to similar trials than repeats, we directly compared the similar and repeat trials with paired sample t-tests in these areas. The difference between similar trials and repeats was significant in both the left PRC (M_similar_ = 0.05, SD_similar_ = 0.13, M_repeat_ = 0.02, SD_repeat_ = 0.14, *t*(38) = 2.51, *p* = .028) and right PRC (M_similar_ = 0.07, SD_similar_ = 0.16, M_repeat_ = 0.03, SD_repeat_ = 0.16, *t*(38) = 2.34, *p* = .037), suggesting that the PRC was sensitive to the object similarity manipulation. However, the difference between similar trials and repeats was not significant in either the left (M_similar_ = 0.02, SD_similar_ = 0.16, M_repeat_ = 0.0, SD_repeat_ = 0.17, *t*(38) = 1.23, *p* = .271) or right CA3DG (M_similar_ = 0.06, SD_similar_ = 0.17, M_repeat_ = 0.04, SD_repeat_ = 0.16, *t*(38) = 1.55, *p* = .257). We observed a strong repetition suppression to repeats relative to first presentation of repeats bilaterally in the PRC (left PRC: M_first_rep_ = 0.07, SD_first_rep_ = 0.14, M_repeat_ = 0.02, SD_repeat_ = 0.14, *t*(38) = 3.26, *p* = .007; right PRC: M_first_rep_ = 0.1, SD_first_rep_ = 0.16, M_repeat_ = 0.03, SD_repeat_ = 0.16, *t*(38) = 3.8, *p* = .002). This comparison did not reveal a significant difference in the CA3DG (left CA3DG: M_first_rep_ = 0.04, SD_first_rep_ = 0.16, M_repeat_ = 0.0, SD_repeat_ = 0.17, *t*(38) = 1.75, *p_unadjusted_*= .089; right CA3DG: M_first_rep_ = 0.06, SD_first_rep_ = 0.16, M_repeat_ = 0.04, SD_repeat_ = 0.16, *t*(38) = 1.78, *p_unadjusted_* = .083).

To directly test whether the PRC was more sensitive to object information in our task than the CA3DG, we ran a repeated measures ANOVA with the within factors presentation rank (first vs. second), trial type (similar vs. repeat), area (PRC vs. CA3DG) and hemisphere. Comparing the PRC and the CA3DG revealed a main effect of presentation rank (*F*(1, 38) = 18.8, *p* < .001), an interaction between presentation rank and area (*F*(1, 38) = 5.3, *p* = .027), and a three-way interaction between presentation rank, area, and hemisphere (*F*(1, 38) = 6.29, *p* = .017). The interaction effect between presentation rank and area expresses that there was a stronger decrease in activity for second presentations (repeats and similar trials) in the PRC than in the CA3DG.

#### Similarity level model

As a follow-up, we binned similar trials in the encoding into high- and low-similarity trials, which enabled us to analyze percent signal change as a function of similarity. For this analysis, we merged all first presentations into one explanatory variable, given that previously we did not find a baseline difference between the first presentations of different types. This resulted in four regressors of interest: Repeat, High-Similarity, Low-Similarity, and New trials. Item similarity level dependent activations from each region that showed a main effect of presentation rank in the previous balanced model were tested with a one-way repeated measures ANOVA. We found a significant effect of similarity in each region included in this analysis: left CA3DG (*F*(3, 114) = 3.35, *p* = .022, *η^2^_partial_* = .08), left SUB (*F*(3, 114) = 3.96, *p* = .01, *η^2^_partial_* = .09), left PRC (*F*(3, 114) = 9, *p* < .001, *η^2^_partial_* = .19), right PRC (*F*(3, 114) = 12.04, *p* < .001, *η^2^_partial_* = .24). Post-hoc comparisons revealed that low similarity trials elicited significantly higher activation than repeats in the bilateral PRC (left PRC: *M_low_sim_* = 0.05, *SD_low_sim_* = 0.14, *M_repeat_* = 0.01, *SD_repeat_* = 0.15, *t*(38) = 2.43, *p* = .03, right PRC: *M_low_sim_* = 0.07, *SD_low_sim_* = 0.17, *M_repeat_* = 0.02, *SD_repeat_* = 0.16, *t*(38) = 3.2, *p* = .004), and high similarity trials elicited marginally higher activation than repeats in the left PRC (*M_high_sim_*= 0.04, *SD_high_sim_*= 0.12, *t*(38) = 1.96, *p* = .068, *p_unadjusted_* = .057). No other areas expressed significant sensitivity to similar trials relative to repeats.

#### Brain-behavioral correlations in the Object Task

To assess whether medial temporal lobe signals were related to participants’ item discrimination performance, we first obtained the participant-specific slopes of sensitivity to item similarity from each region that showed any modulation by the Object Encoding conditions (left CA3DG, left SUB, left and right PRC). We achieved this by building linear mixed effects models, predicting each region’s activity based on the level of similarity (Repeat, High Similarity, Low Similarity, New) with random slopes and random intercepts, and then extracted the participant-specific coefficients. Likewise, we used participant specific slopes of behavioral sensitivity to item similarity, obtained from linear mixed models predicting “new” responses based on the two similarity variables (between the first encoding presentation and recognition, and the second encoding presentation and recognition). We used Spearman correlations to assess the relationship between the neural and behavioral measures of item similarity sensitivity, and controlled for the FDR as before. Sensitivity of the right PRC correlated significantly with participants’ behavioral sensitivity to the similarity between the second encoding trial and the recognition trial (*r* = 0.41, *p* = .035), indicating that the steeper the response of the right PRC was to increasing levels of similarity, the higher behavioral sensitivity the participant expressed to the similarity manipulation, which manifested as the probability of giving a “new” response to lure and foil fractals. Additionally, we found a marginally significant negative correlation between the sensitivity of the left CA3DG to object similarity and behavioral sensitivity to the second similarity measure, but this association was not significant after adjusting for multiple comparisons (*r* = −0.31, *p* = .1, *p_unadjusted_* = .051). No other ROI revealed any significant correlation between behavioral and neural sensitivity to similarity (all *r*s < |.2|, *p*s > .1).

### Location encoding: Univariate ROI results

#### Presentation rank by trial type model

In the Location Task, a region-wise 2-by-2 repeated measure ANOVA of presentation rank (first and second) and trial type (similar and repeat) revealed a main effect of presentation rank in the following regions: left CA3DG (*F*(1, 38) = 8.53, *p* = .012, *η^2^_partial_* = .18), right CA3DG (*F*(1, 38) = 11.01, *p* = .008, *η^2^_partial_* = .22), right SUB (*F*(1, 38) = 10.85, *p* = .008, *η^2^_partial_* = 0.22), left PHC (*F*(1, 38) = 9.71, *p* = .009, *η^2^_partial_* = 0.2), and right PHC (*F*(1, 38) = 10.47, *p* = .008, *η^2^_partial_* = 0.22). The effect in the left CA1 was not significant after correcting for multiple comparisons (*F*(1, 38) = 4.31, *p* = .074, *p_unadjusted_* = .045, *η^2^_partial_* = .1). The main effect of presentation rank was driven by a lower signal at Presentation 2 indicating wide-spread repetition suppression. In contrast, we found no effect in either the left or the right PRC (both *p*s > .1), suggesting that this area did not show repetition suppression during Location Encoding. We did not find a main effect of trial type (all *p*s > .2), or any significant difference between first presentation of similar trials and first presentation of repeat trials in any area (all *p*s > .2), and thus we concluded that there was no baseline difference between the first presentations of different types. Of note, the interaction between presentation rank and trial type in the left PHC was marginally significant (*F*(1, 38) = 3.83, *p_unadjusted_*= .058, *η^2^_partial_* = 0.092, Figure 4c). Post-hoc paired sample t-test in the left PHC revealed that while activity was significantly higher for first presentations of later repeats than repeats (M_first_rep_ = 0.06, SD_first_rep_ = 0.14, M_repeat_ = 0.01, SD_repeat_ = 0.1, *t*(38) = 2.8, *p* = .024), there was no significant difference between first presentations of later similar trials and similar trials (M_first_sim_ = 0.05, SD_first_sim_ = 0.12, M_similar_ = 0.04, SD_similar_ = 0.12, *t*(38) = 1.31, *p* = .236). This pattern was also found in the right PHC (first presentation of later repeats vs. repeats: M_first_rep_ = 0.09, SD_first_rep_ = 0.14, M_repeat_ = 0.04, SD_repeat_ = 0.11, *t*(38) = 2.87, *p* = .022, first presentations of later similar trials vs. similar trials: M_first_sim_ = 0.08, SD_first_sim_ = 0.13, M_similar_ = 0.06, SD_similar_ = 0.12, *t*(38) = 1.64, *p* = .163). Activity was additionally significantly higher for similar trials than repeats in the left PHC, but this effect did not persist after adjusting for multiple comparisons (*t*(38) = 2.22, *p* = .065, *p_unadjusted_* = .033). We expected that the activation to similar trials would be higher than to repeats in the CA3DG, given its hypothesized role in pattern separation, however, the difference was not significant in either the left (M_similar_ = 0.03, SD_similar_ = 0.14, M_repeat_ = 0.05, SD_repeat_ = 0.14, *t*(38) = −1.17, *p* = .301) or right CA3DG (M_similar_ = 0.04, SD_similar_ = 0.15, M_repeat_ = 0.03, SD_repeat_ = 0.14, *t*(38) = 0.4, *p* = .69). We found repetition suppression to repeats relative to the first presentation of repeats in the right CA3DG, which was not significant after adjusting for the FDR (M_first_rep_ = 0.06, SD_first_rep_ = 0.15, M_repeat_ = 0.03, SD_repeat_ = 0.14, *t*(38) = 2.06, *p* = .092, *p_unadjusted_* = .046). Note, however, that the main effect of presentation rank in the bilateral CA3DG nonetheless indicates that the activity to similar trials and repeats was lower than to first presentations in this region.

To directly test whether the PHC was more sensitive to the location information in our task than CA3DG, we ran a repeated measures ANOVA with the within factors presentation rank (First vs. Second), trial type (Similar vs. Repeat), area (PHC vs. CA3DG) and hemisphere. Comparing the PHC and the CA3DG revealed a main effect of presentation rank (*F*(1, 38) = 18.7, *p* < .001) and a three-way interaction between presentation rank, trial type, and area (*F*(1, 38) = 5.51, *p* = .024).

**Figure 6.**
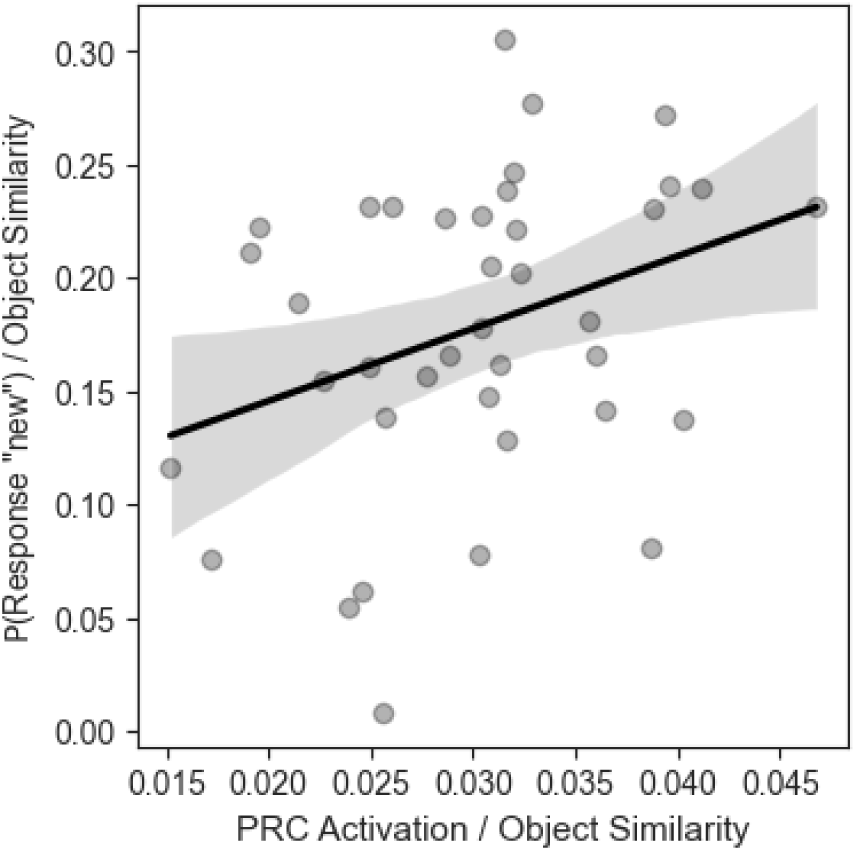
The association between the right PRC signal and mnemonic discrimination in the object domain. The behavioral sensitivity to object similarity - operationalized as the individual slope of similarity dependent “new” responses - correlates with the steepness of the right PRC’s response to the level of object similarity.

#### Similarity level model

We binned similar trials of the Location Encoding into high- and low-similarity trials to analyze percent signal change as a function of spatial similarity. Again, all first presentation trials were merged into one regressor, creating four similarity regressors: Repeat, High-Similarity, Low-Similarity, and New trials. In the statistical analysis of the extracted percent signal change values, we included all regions of interest that showed a response to the task manipulation in the previous analysis, i.e. left CA1, left and right CA3DG, right SUB, left PHC, and right PHC. Spatial similarity level dependent activations from each of these regions are shown in Figure 7. We assessed the sensitivity to spatial similarity in a one-way repeated measures ANOVA. We found a main effect of similarity in all regions included in this analysis (left CA3DG: *F*(3, 114) = 4.24, *p* = .007, *η^2^_partial_* = .1, right CA3DG: *F*(3, 114) = 2.98, *p* = .034, *η^2^_partial_* = .07, right SUB: *F*(3, 114) = 3.17, *p* = .033, *η^2^_partial_* = .07, left PHC: *F*(3, 114) = 3.98, *p* = .01, *η^2^_partial_* = .09, right PHC: *F*(3, 114) = 2.93, *p* = .037, *η^2^_partial_* = .07), except the left CA1 (F(3, 114) = 2.17, p = .094, *η^2^_partial_* = .05). Post-hoc comparisons revealed that the left PHC was the only area where the difference between repeated and highly similar trials was significant, albeit this difference did not persist after adjusting for multiple comparisons (*M_high_sim_*= 0.04, *SD_high_sim_*= 0.11, *M_repeat_*= 0.01, *SD_repeat_*= 0.1, *t*(39) = 2.19, *p* = .104, *p_unadjusted_* = .035).

**Figure 7.**
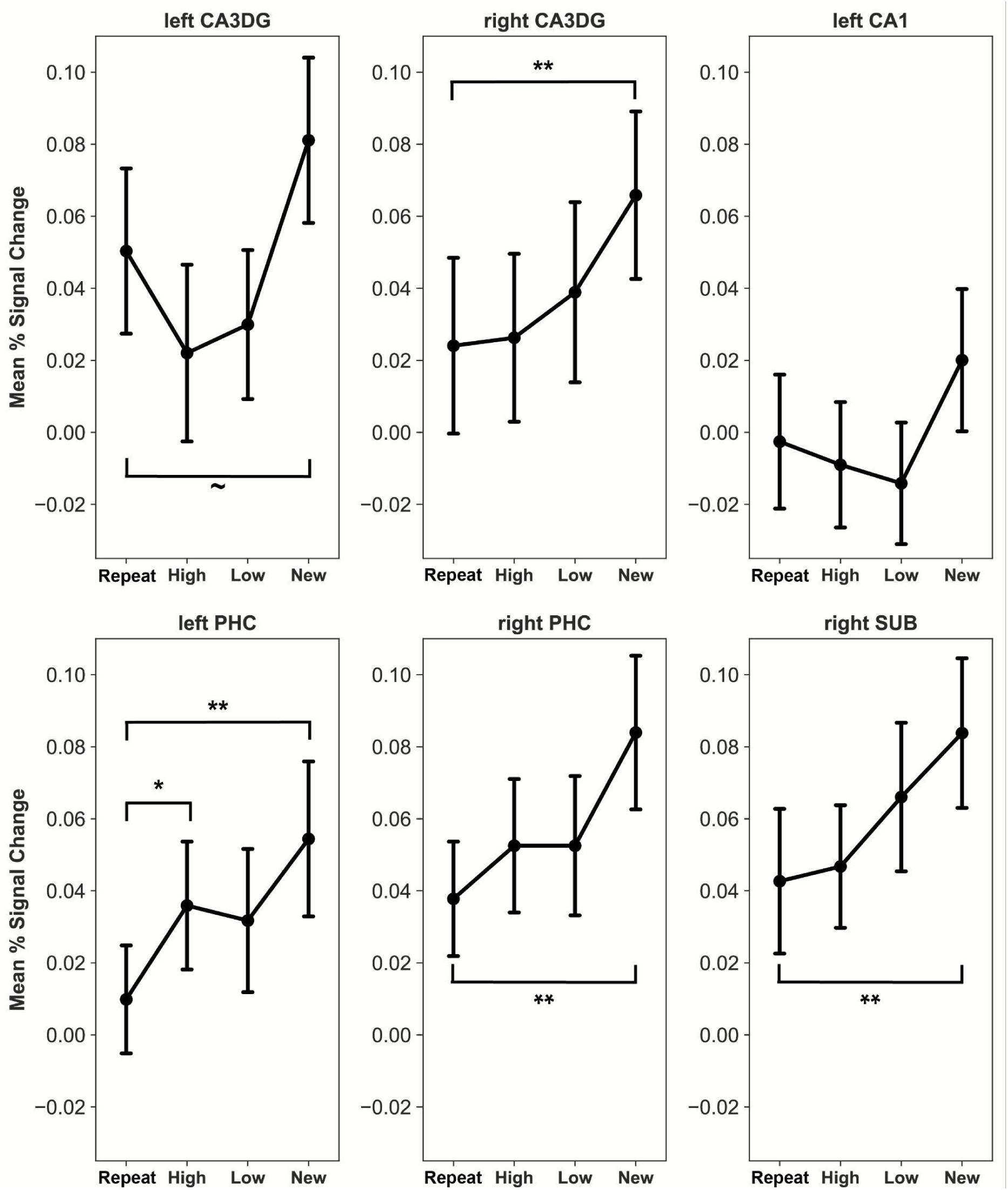
Similarity dependent activations during location encoding. The left PHC expresses a significant increase to trials with high similarity. Significant differences relative to repeats (unadjusted) are denoted with a symbol: ∼ p < .06, * p < .05, ** p < .01. Responses from all areas that showed a significant effect in the previous 2-by-2 model were included in the figure. Abbreviations: CA1 - cornu Ammonis 1 and 2, CA3DG - cornu Ammonis 3 and dentate gyrus, SUB - subiculum,, PHC - parahippocampal cortex. Error bars represent the standard error of mean.

#### Brain behavioral correlations in the Location Task

We then assessed the associations between the neural responses of the medial temporal ROIs and mnemonic discrimination in the spatial domain. For this purpose, we obtained the participant-specific slopes of sensitivity to spatial similarity from linear mixed effects models that predicted each region’s responses based on the level of similarity. All regions that exhibited some modulation by the Location Encoding conditions were included in this analysis (left CA1, left and right CA3DG, right SUB, left PHC, and right PHC). Each participant’s behavioral sensitivity (i.e. individual slope) to the spatial similarity between the first encoding presentation and the recognition trial, and the second encoding presentation and the recognition trial was used as a behavioral outcome. We ran Spearman correlations, and found an association between the right CA3DG response and the behavioral sensitivity to the similarity between the second encoding trial and the recognition trial (*r* = 0.34, *p_unadjusted_* = .034, Supplementary Figure 1) that did not remain significant after adjusting for the FDR (*p* = .168).

### CA3DG responses combined across tasks

The results above indicated that the PRC and PHC showed more pronounced responses than the CA3DG during the Object and Location encoding, respectively, and that the CA3DG showed little sensitivity to similar trials. As a follow-up analysis, we combined the extracted percent signal change values of the CA3DG from the two tasks, and conducted one repeated measures ANOVA per hemisphere with similarity level (repeat, high similarity, low similarity, new) and task (Object versus Location) as within-subject factors. In both the left and right CA3DG, we found a main effect of similarity level (left CA3DG: *F*(3, 114) = 8.18, *p* < .001, right CA3DG: *F*(3, 114) = 6.13, *p* < .001), but no effect of task and no task-similarity interaction (all *p*s > 0.3). In the left CA3DG post-hoc paired sample t-tests between similarity levels revealed a significant difference between repeat and new trials (*t*(77) = 3.32, *p* = .004), high similarity trials and new trails (*t*(77) = 4.22, *p* < .001), and low similarity trials and new trials (*t*(77) = 3.07, *p* < .006). There was no significant difference between either high or low similarity and repeat trials (all *p*s > 0.3). In the right CA3DG post-hoc paired sample t-tests between similarity levels revealed a significant difference between repeat and new trials (*t*(77) = 3.95, *p* = .001), and high similarity trials and new trails (*t*(77) = 3.16, *p* = .007). There was no significant difference between high similarity and repeat trials (*p* > 0.5). The difference between the Low similarity and repeat trials was marginally significant without adjustment (*t*(77) = 1.91, *p* = .118, *p_unadjusted_* = .059). Further, low similarity trials did not significantly differ from new trials (*t*(77) = 1.08, *p_unadjusted_* = .285). Altogether, these results are in line with the previous task-wise analysis, i.e., that the CA3DG shows repetition suppression to second presentation trials, and although there seems to be some similarity based modulation in the CA3DG, the difference between repeated and similar trials is not statistically significant.

### Task-dependent responses in the parahippocampal and perirhinal cortex

We tested whether the activity of PRC and PHC statistically differed depending on the task context, i.e., whether the PRC is more engaged in object interference resolution than the PHC and vice versa. To this end, we merged data from the two tasks and two regions, and conducted a repeated measures ANOVA with area (PHC versus PRC), hemisphere, similarity level (Repeat, High Similarity, Low Similarity, New), and task (Object versus Location) as within-subject factors. We found a significant main effect of similarity level (*F*(3, 114) = 13.07, *p* < .001), and a main effect of hemisphere (*F*(1, 38) = 13.07, *p* = .043). The main effect of hemisphere was due to higher mean percent signal change in the right hemisphere (*t*(623) = 4.08, *p* < .001). We did not find a main effect of either task (*F*(1, 38) = 0.01, *p* = .922) or area (*F*(1, 38) = 1.28, *p* = .266). Importantly, the analysis revealed a significant three-way interaction between similarity level, area, and task (*F*(3, 114) = 3.87, *p* = .011). This interaction indicates that depending on the task context – and thus the type of information that needed to be attended – similarity induced interference differentially modulated the activity of the PRC and PHC (Figure 8).

**Figure 8.**
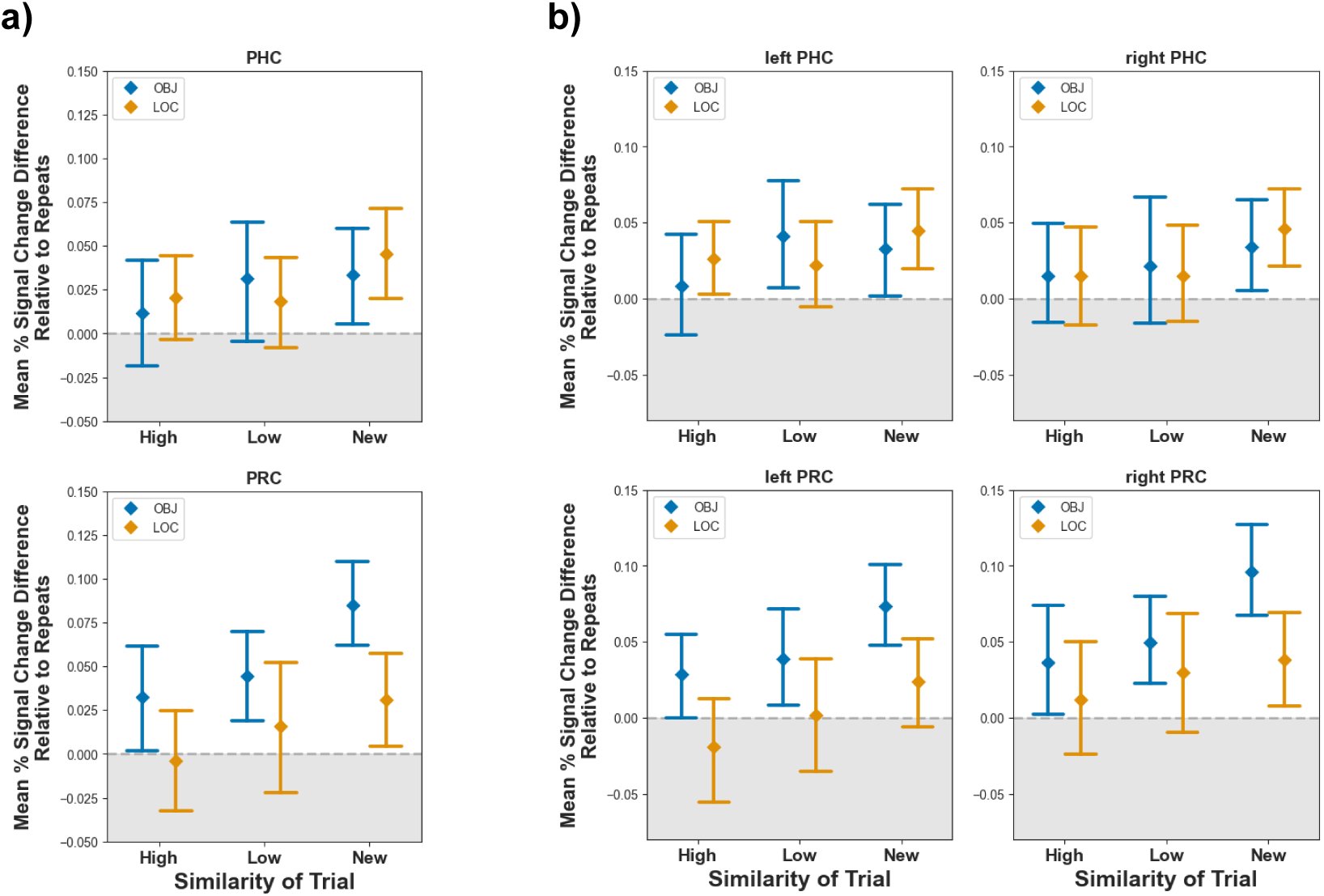
Similarity level dependent responses of the parahippocampal and perirhinal cortices across the two tasks. Responses to repeats were extracted from the mean percent signal change values of each similarity level for visualization purposes. Values around and below 0 therefore can be considered repetition suppression. Panel a) shows the cross-hemisphere average, and panel b) shows the signal from each hemisphere. A three-way interaction between similarity-level, task and area indicates that similarity dependent responses differ based on task context. Error bars represent 95% confidence intervals.

### Whole-brain results

#### Univariate activations

We found 13 clusters in the Object Encoding (Figure 9a) and 12 clusters in the Location Encoding (Figure 9b) that showed significantly higher activity to similar trials than their first presentation, summarized in Supplementary Table 6 and 7. Importantly, the clusters derived from the Location and Object task moderately overlapped (Dice-Sørensen coefficient = 0.54, Figure 9c), and both covered regions belonging to the frontoparietal control network, i.e., the middle frontal gyrus and the supramarginal gyrus (Dixon et al., 2018; Uddin et al., 2019) and the angular gyrus. No significant clusters were found in the medial temporal lobe. Results from the contrast (*similar - first presentation of later similar trials) > (repeats - first presentation of later repeats)* are shown in Figure 10 and reported in Supplementary Table 4 and 5. Untresholded z maps were uploaded to Neurovault (Gorgolewski et al., 2016) and analyzed with Neurosynth (Yarkoni et al., 2011) to obtain the most closely associated neurocognitive terms based on the meta-analysis of the available fMRI data. This analysis supported our interpretation that frontoparietal regions were activated by similar trials. In turn, the terms “default mode”, “memory retrieval”, “recognition” and “episodic memory” were less associated with the similar > first activation maps. Results of the Neurosynth analysis (including the top 10 associated terms) for the similar > first contrast are reported in the Supplementary Tables 8-11.

**Figure 9.**
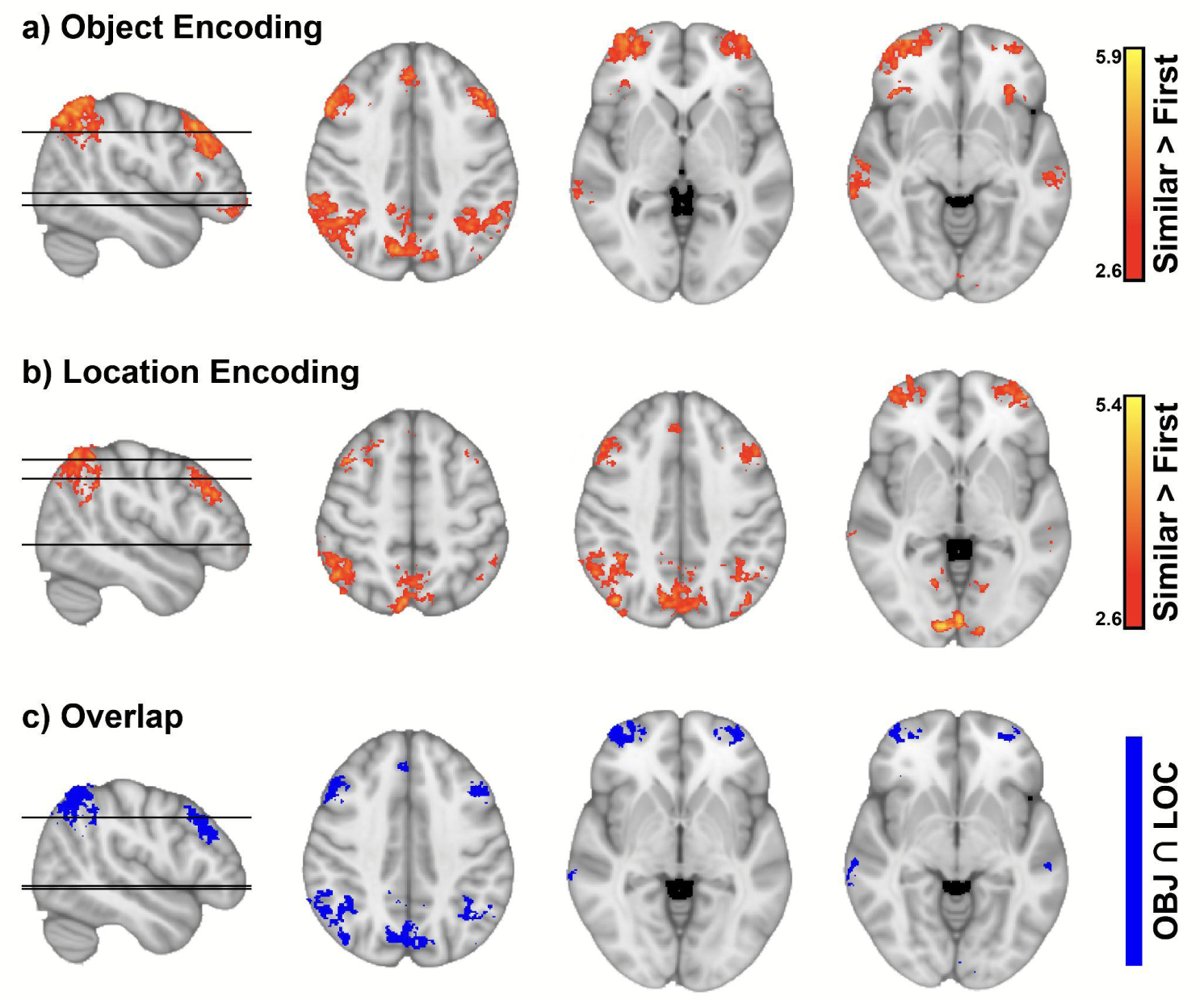
Clusters of significant activation to similar > first trials. Whole brain activations to similar > first presentation trials showed an overlap between the Object and Location Tasks. Frontoparietal regions were activated by both task contexts. a) Significant clusters of activation during the Object Encoding. b) Significant clusters of activation during the Location Encoding. c) Overlap of significant clusters in the Location and Object Tasks.

**Figure 10.**
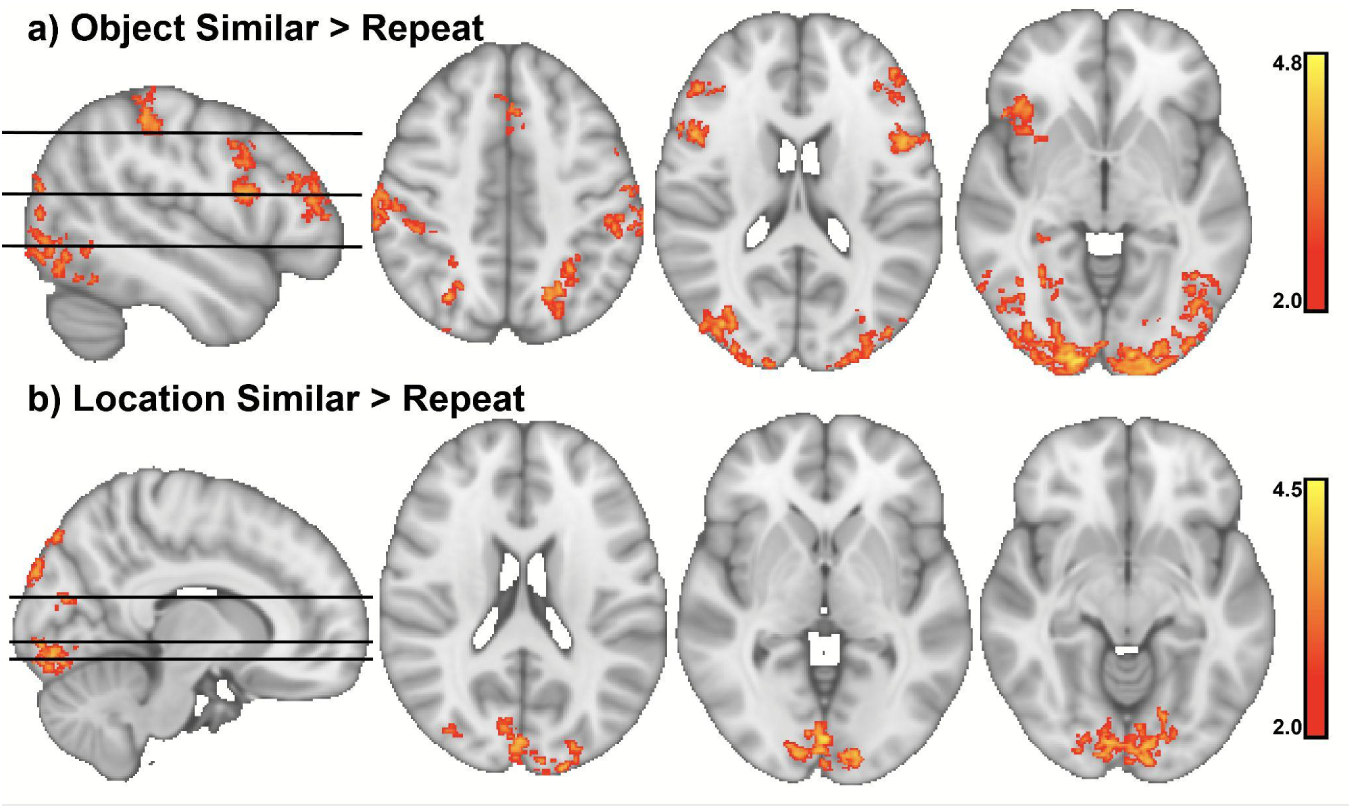
Clusters of significant difference between similar and repeat trials. The figure depicts the contrast (similar > first presentation) > (repeat > first presentation), which isolates regions specifically activated during the encoding of similar stimuli. a) During similar object trials, occipital areas, as well as the superior parietal gyrus, the inferior frontal gyrus, and middle frontal gyrus were activated more than during repeat trials. b) During similar location trials, occipital areas were activated more so than during repeat trials.

#### gPPI with frontal and parietal seeds

We report results from each seed region’s interaction with the similar (similar PPI contrast) and repeat condition (repeat PPI contrast), which reflect increases in connectivity relative to the baseline. Cluster statistics for the similar PPI contrast can be found in Table 1 (Object Task) and Table 2 (Location Task). Additionally, we report results from the contrast similar PPI > repeat PPI, which indicate if the seed region interacts with some areas more in the similar condition than the repeat condition. One gPPI analysis was run per ROI and task. In the Object Task, only the right angular gyrus and the right middle frontal cortex seeds expressed increased functional connectivity with other regions. We found 1 cluster (256 voxels, max. *z* = 3.42, *p* = .044) of increased connectivity during the encoding of similar object trials between the right angular gyrus seed and the left supramarginal gyrus and left angular gyrus. The right middle frontal gyrus seed increased its connectivity with two clusters: one located in the left angular cortex (301 voxels, *z* = 3.5, *p* = .011), and the other in the posterior region of the middle temporal gyrus (343 voxels, *z* = 3.64, *p* = .004). No significant clusters were found for the contrast similar PPI > repeat PPI in the Object Task. This indicates that the interaction between similar trials and functional connectivity did not statistically differ from the interaction between repeat trials and functional connectivity. Note, however, that no significant clusters were found for the repeat PPI contrast in the Object task.

**Table 1.** Significant clusters in the Object Task for the similar PPI contrast.

| Seed region | Areas in cluster | N. voxels | Max. z | Cluster p |
| --- | --- | --- | --- | --- |
| right angular gyrus | left supramarginal gyrus,<br>left angular gyrus | 256 | 3.42 | .044 |
| right middle frontal gyrus | left angular gyrus | 301 | 3.5 | .011 |
|  | left middle temporal gyrus | 343 | 3.63 | .004 |

**Table 2.** Significant clusters in the Location Task for the similar PPI contrast.

| Seed region | Areas in cluster | N. voxels | Max. z | Cluster p |
| --- | --- | --- | --- | --- |
| **left angular gyrus** | right angular gyrus and<br>supramarginal gyrus | 383 | 3.33 | .002 |
|  | right frontal pole | 373 | 2.92 | .002 |
|  | right postcentral gyrus and<br>supramarginal gyrus | 282 | 3.05 | .023 |
|  | right supramarginal gyrus | 266 | 3.82 | .037 |
| **right angular gyrus** | left angular gyrus,<br>left supramarginal gyrus | 256 | 3.42 | .044 |
| **left supramarginal gyrus** | right angular gyrus | 2055 | 3.69 | < .001 |
|  | right frontal pole | 1141 | 3.62 | < .001 |
|  | right middle temporal gyrus, posterior division | 931 | 3.91 | < .001 |
|  | right posterior cingulate and precuneus | 899 | 3.3 | < .001 |
|  | left lateral occipital cortex | 426 | 3.44 | < .001 |
|  | left precuneus | 296 | 3.3 | .013 |
|  | right superior frontal gyrus and frontal pole | 259 | 3.55 | .037 |
| **right supramarginal gyrus** | right angular gyrus and lateral occipital cortex | 3016 | 3.68 | < .001 |
|  | left frontal pole | 1607 | 3.56 | < .001 |
|  | right middle temporal gyrus, posterior division | 1455 | 3.47 | < .001 |
|  | bilateral lingual gyrus | 1300 | 3.81 | < .001 |
|  | right frontal pole | 1087 | 3.99 | < .001 |
|  | left inferior and middle frontal gyrus | 1056 | 3.72 | < .001 |
|  | right middle frontal gyrus | 845 | 4 | < .001 |
|  | left middle frontal gyrus | 589 | 4.14 | < .001 |
|  | right middle frontal gyrus, precentral gyrus | 549 | 3.1 | < .001 |
|  | bilateral precuneus | 476 | 3.2 | < .001 |
|  | right fusiform gyrus | 426 | 3.37 | < .001 |
|  | bilateral paracingulate gyrus | 413 | 3.55 | < .001 |
|  | right superior frontal gyrus | 405 | 3.33 | < .001 |
|  | bilateral posterior cingulate gyrus | 402 | 3.43 | < .001 |
|  | left lingual gyrus, left fusiform gyrus | 372 | 3.54 | .002 |
|  | left paracingulate gyrus, left anterior cingulate gyrus | 355 | 3.39 | .002 |
|  | left lateral occipital cortex, left angular gyrus | 350 | 3.59 | .003 |
|  | left orbitofrontal cortex | 331 | 3.44 | .005 |
|  | right lateral occipital cortex | 315 | 3.36 | .007 |
|  | right middle temporal gyrus, anterior division | 304 | 3.13 | .01 |
| **left middle frontal gyrus** | right frontal pole | 467 | 3.09 | < .001 |
|  | right angular gyrus, right supramarginal gyrus | 328 | 3.41 | .005 |

**Table 3.**
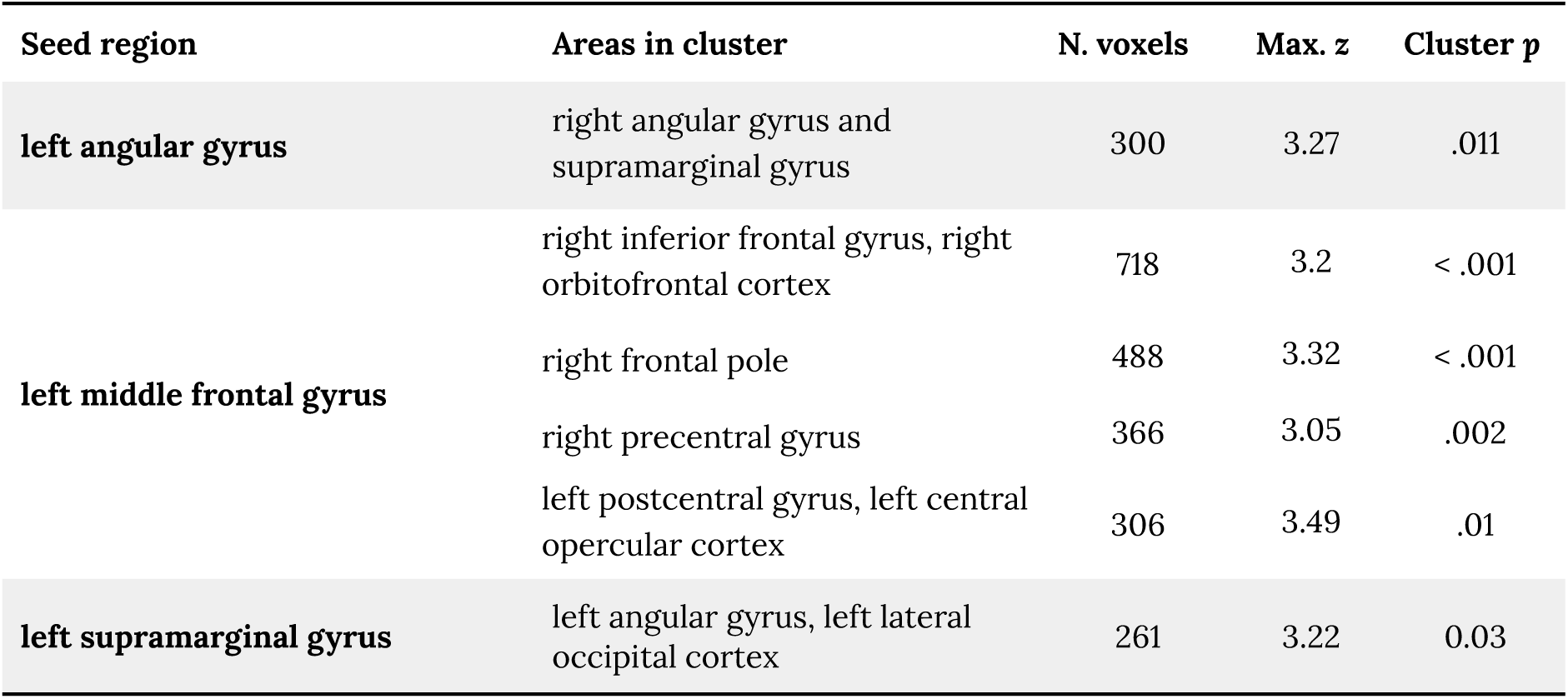
Significant clusters in the Location Task for the repeat PPI contrast.

In the Location Task, we found enhanced functional connectivity between the left angular gyrus seed and the right frontal pole, right supramarginal gyrus, right postcentral gyrus, and right angular gyrus during the encoding of similar spatial trials. The left supramarginal gyrus increased its connectivity with the right frontal pole and superior frontal gyrus, right middle temporal gyrus, bilateral cingulate and precuneus cortex, right angular and supramarginal gyrus, and left lateral occipital cortex. The right supramarginal gyrus expressed a similar, but even more extensive pattern of connectivity increase, involving some additional sensory areas, such as the lingual gyrus and occipital fusiform cortex, in addition to bilateral frontal and parietal regions. As for the left middle frontal gyrus seed, only two significant clusters of connectivity increase were found, one in the right frontal pole and one in the right angular gyrus. No significant connectivity increase was found for the right angular gyrus and right middle frontal gyrus seeds for the similar PPI contrast.

Moreover, we found two significant, anatomically overlapping clusters for the similar PPI > repeat PPI contrast in the Location task. Both the left supramarginal gyrus (363 voxels, max. *z* = 3.79, *p* = .001) and the right middle frontal gyrus (23 voxels, max. *z* = 3.5, *p* = .006) increased their connectivity with the left occipital pole during the encoding of similar trials, and this increase was significantly higher than that in the repeat condition.

### Frontoparietal-medial temporal functional connectivity

Each medial temporal ROI exhibited significant connectivity with the frontoparietal regions in both tasks (Supplementary Figure 3 and 4). Further, the connectivity between the right PHC and the left supramarginal gyrus was significantly higher during the Location Encoding than the Object Encoding (*t*(39) = 2.23, *p* = .025), but this difference was not significant after adjusting for the FDR (*p* = .521). No other difference was found in the connectivity of medial temporal and frontoparietal regions between the two tasks (Figure 11).

**Figure 11.**
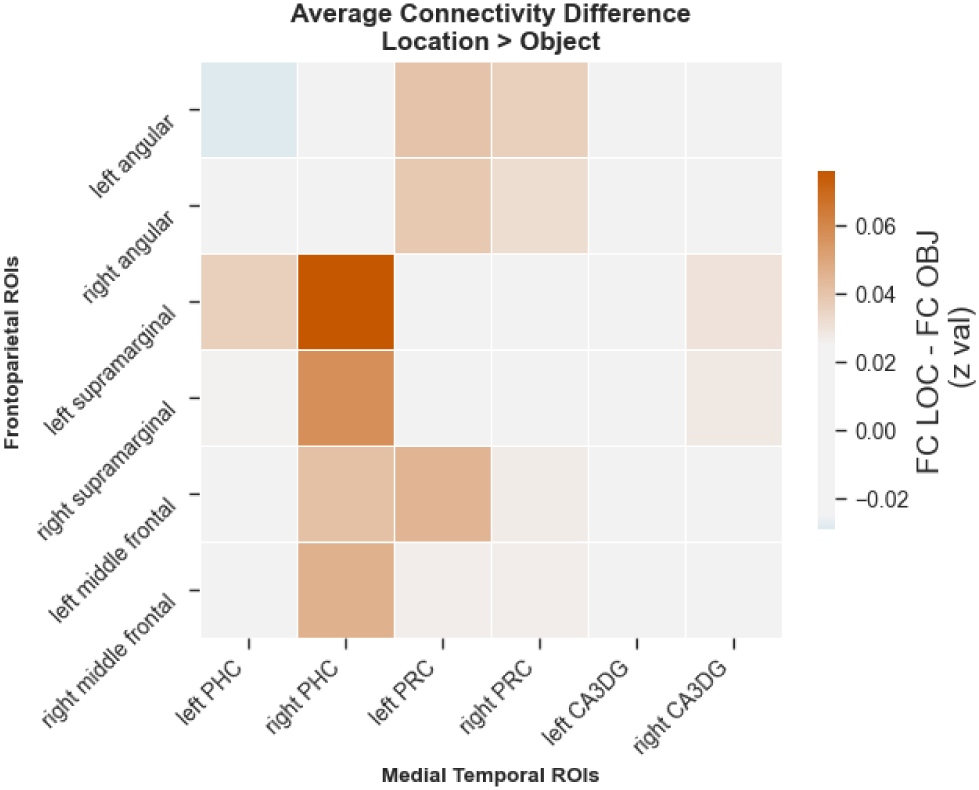
Functional connectivity differences between the Object and the Location Encoding. The connectivity between the right PHC and left supramarginal gyrus was higher during Location Encoding than Object Encoding. The correlation between each medial temporal ROI and each frontoparietal ROI during the Object Encoding was subtracted from the Location Encoding for the purpose of visualizing the differences in connectivity between the tasks.

We then proceeded to assess the associations between the medial temporal - frontoparietal connectivity and behavior. We correlated the behavioral variables of object and spatial similarity sensitivity with the connectivity between each medial temporal ROI and frontoparietal ROI. Intriguingly, we found exclusively negative relationships between the connectivity of both the PHC and PRC during encoding and the behavioral sensitivity to similarity. Due to the number of comparisons (36 connectivity variables), none of the correlations remained significant after FDR adjustment. We report these associations in order of effect size in the supplementary. Of interest, in the Object task the sensitivity to the similarity between the first encoding presentation and the recognition trial was associated with the connectivity between the right PRC and right angular gyrus (*r* = −0.41, *p* = .01). The sensitivity to the similarity between the second encoding presentation and the recognition trial was associated with the connectivity between the right PHC and left supramarginal gyrus (*r* = −0.41, *p* = .009), and the left PRC and right supramarginal gyrus (*r* = −0.41, *p* = .009). In the Location task the associations were weaker, but also exclusively negative, and included both the PRC and PHC connectivity (see Supplementary Table 13). We found no associations between the behavioral measures and the connectivity of the CA3DG in either task.

## Discussion

Our results provide three contributions to the cognitive neuroscience of memory. First, amid the debate about the anatomical and functional independence of the two MTL pathways (Burke et al., 2018; Connor & Knierim, 2017; Lawrence et al., 2020; Nilssen et al., 2019; Rolls, 2024), our results support the notion that the perirhinal and parahippocampal cortices provide complementary information about individual items and their spatial context, respectively. Second, we demonstrate that the perirhinal and parahippocampal cortices show a sensitivity to interfering mnemonic information, even when this information is not easily verbalized and categorized. In contrast, we found only limited evidence that responses were modulated by the similarity of non-meaningful stimuli in the CA3DG. Third, our results suggest that frontoparietal regions are recruited in response to both spatial and object interference, and the connectivity of these regions flexibly changes according to task demands. Below, we discuss the interpretation of these results, and place them within a general discussion on medial temporal lobe function and human memory.

Behaviorally, we found a significant difference between the ratio of “new” responses to target and lure trials in both the Object and Location Tasks. Moreover, we found that not only the first, but the similarity of the second encoding trial influenced participants’ discrimination behavior during recognition. Together, these behavioral results strongly suggest that mnemonic discrimination was affected by the similarity manipulation at encoding. Therefore, we can conclude that the interference between the encoding trials provides a viable basis for assessing pattern separation at the neural level. A noteworthy difference between the Object and Location tasks is the inclusion of foils (unique, never-before-seen items) in the Object recognition, whereas only lures and distant lures were included in the Location recognition. Our goal was to make the two tasks as similar as possible, but due to the difference in the type of information that was manipulated - namely, that the location information has to bind to a specific object - some asymmetry in design was unavoidable. All neuroimaging results reported here are based on the encoding phase of the tasks, and thus they are not affected by this difference during recognition.

In line with prior fMRI work (Bencze et al., 2024; Berron et al., 2018; Reagh & Yassa, 2014), our results support the view that a division of labor exists between the PRC and PHC. The PHC showed strong repetition suppression to repeated trials viewed in unchanged locations, and we found some tentative evidence that its activity was modulated by high-similarity spatial trials. Conversely, the experimental manipulations of object features elicited significant effects in the PRC, including repetition suppression to repeated trials, and activity changes as a function of similarity. A direct comparison of the effects of similarity on the PRC and PHC in the Object and Location Task revealed a three-way interaction of similarity, area, and task, demonstrating that the responses of these areas differed based on task context. Notably, new and repeated trials had an identical structure during location and object encoding, yet the PRC showed sensitivity to the task context, and expressed stronger repetition suppression to repeated trials when the task demands matched its respective specialization. This highlights the possibility that PRC and PHC responses are modulated by top-down control processes rather than pure trial features.

The PRC and PHC showed sensitivity to highly similarity trials in accordance with their task domains. Specifically, in the Object Task we found that similar trials elicited significantly higher activation than repeated trials in both the left and right PRC. Moreover, the sensitivity of the PRC to object similarity was related to participants’ mnemonic discrimination behavior. Correspondingly, in the Location Task we found a tendency of high-similarity trials to elicit higher activation than repeated trials in the left PHC. When considered together with the result that the activation to similar trials did not differ statistically from first presentation trials in either the left or and right PHC, this suggests that the PHC was sensitive to interfering location information. On the other hand, the CA3DG did not express any significant increase in response to similar trials in either task or when responses of the two tasks were combined. Additionally, directly comparing the CA3DG with the PRC and PHC in the Object and Location task revealed that the neocortical regions showed higher task sensitivity in their respective domains than the CA3DG. Specifically, we found an interaction between area (CA3DG vs. PRC) and presentation rank in the Object task, meaning that the PRC showed a more pronounced decrease to second presentation trials than the CA3DG, i.e., repetition suppression to fractals was stronger in the PRC. In the Location task, a three-way interaction between presentation rank (first vs. second), trial type (repeat vs. similar), and area (PHC vs. CA3DG) suggests that PHC responses were modulated more by repeats and similar trials than those of the CA3DG. These results indicate that the activity of neocortical areas in the medial temporal lobe may be sensitive to high-interference trials even when the CA3DG exhibits only moderate activity increase in response to similarity, suggesting that the hippocampus is less involved in the differentiation of non-meaningful stimuli than the neocortex.

Numerous previous studies suggest the dentate gyrus is involved in creating unique memories in humans (e.g., Azab et al., 2014; Bakker et al., 2008; Berron et al., 2016; Kyle et al., 2015), and some have argued for a necessary role of the dentate gyrus in mnemonic discrimination (Baker et al., 2016; Mitchnick et al., 2024). A key difference between preceding work and ours is the meaningfulness of stimuli used. A few studies investigating pattern separation used non-meaningful stimuli, and found evidence of hippocampal involvement, but these either included items that resembled real objects and were verbally labeled (Duff et al., 2012), or required only perceptual (Mitchnick et al., 2022) or working memory discrimination (Paleja et al., 2014). Recently, Wammes and colleagues (2021) used fractals in a statistical learning paradigm and demonstrated that representational change of items with differing levels of similarity corresponds to a U-shaped function in the dentate gyrus. A U-shaped function would be predicted by the nonmonotonic plasticity hypothesis, so that the representation of only moderately similar items (but not low- or high-similarity items) were driven towards differentiation after learning relative to the pre-learning phase. This discrepancy between the results may be explained by the fact that in the study by Wammes et al., 2021 a relatively small set of fractals (16) were presented over a long sequence learning period, thus allowing participants to familiarize themselves well with each individual item. In our study, each similar, interference inducing fractal was presented only once, and their hippocampal representation could not be stabilized through learning. It would be interesting to analyze the data of Wammes et al., 2021, from the beginning of the learning period, which would yield a closer correspondence between the two designs. To the best of our knowledge, this is the first study to examine how interfering, non-meaningful stimuli are encoded in long-term, episodic-like memory through one-shot learning by the CA3DG (cf. Brickman et al., 2014).

Our results align with a representational account of the medial temporal lobe, i.e., the view that areas are not specialized in terms of the type of computation they carry out, but rather according to the type of information they process (Cowell et al., 2019; Martin & Barense, 2023). Advocates of this account see a continuity between the visual and memory systems (Barense et al., 2005; Martin & Barense, 2023; Yonelinas, 2013), with the complexity of representable information gradually increasing from the early visual cortex to the hippocampus. Speculatively, it may be that the fractals did not elicit pattern separation in the hippocampus as images of everyday objects normally do because these stimuli were unfamiliar to participants and thus were comparably lacking in contextual and meaningful features. One interpretation of our results is that only experiences that differ in meaningful features are encoded in long term memory by the CA3DG in a pattern separated fashion. To fully test this hypothesis, the encoding of meaningful and non-meaningful stimuli in the hippocampus should be tested in a single mnemonic discrimination paradigm. Previously literature indicates that meaningful stimuli (i.e., faces and objects) elicit higher activation in the hippocampus relative to novel ones (greebles) in a visual discrimination paradigm (Barense et al., 2011). Future studies could also shed light on whether making participants more familiar with non-meaningful stimuli (e.g., like Wammes et al., 2021), or assigning meaning and associations to them, may facilitate pattern separation for long-term memory in the hippocampus.

Outside of the MTL, we found evidence that the middle frontal gyrus and parietal regions are activated during the encoding of similar trials irrespective of information domain, and their connectivity increases within the network and with downstream sensory regions, when interference resolution is necessary. We interpret this as a regulation of regions that process sensory information in order to either emphasize or downplay differences. These data alone cannot disambiguate, however, whether the connectivity increase facilitated the integration or differentiation of interfering information in this context. A subsequent memory analysis could shed light on how the activation of these regions contributed to memory performance. Our design did not allow for this type of analysis, because each recognition trial can be paired with two encoding trials. However, our behavioral results do show that both the similarity of the first and second presentation encoding trial affected participants recognition performance, indicating that participants did not only “encounter” similar trials, but encoded them. Thus, the activation of frontal and parietal regions during similar trials suggests their involvement in encoding interfering stimuli.

The whole-brain gPPI analysis did not provide evidence that frontal and parietal connectivity with the MTL changed during the encoding of similar trials. This could be related to either the relatively low power of PPI methods for fMRI data (Di & Biswal, 2017; O’Reilly et al., 2012), the lower signal-to-noise ratio and small volume of MTL regions, or a stable connectivity profile between the MTL and the frontoparietal areas. It is important that the direction and amount of influence of the frontoparietal regions may not be directly translatable into changes in functional connectivity, and influence could also be exerted over stable connectivity, which led us to examine the region-to-region connectivity specifically between the MTL and frontoparietal ROIs.

Our region-to-region connectivity analysis between MTL and frontal and parietal seeds confirmed functional connectivity between these networks. Further, we found tentative evidence that the connectivity between the PHC changed according to task context, as the right PHC displayed higher connectivity with the left supramarginal gyrus during location encoding than object encoding. However, no other differences were found in the connectivity patterns of the two tasks. The associations between the MTL-frontoparietal connectivity and recognition performance suggest that the synchronicity of these areas carries behavioral relevance. Previously, suppression of competing memory patterns in ventral visual areas by the lateral prefrontal cortex has been shown to lead to adaptive forgetting of interfering information (Wimber et al., 2015), and numerous studies demonstrated the role of the dorsolateral prefrontal cortex in reducing interference by retrieval suppression and inhibitory control (Anderson et al., 2004; Anderson & Hulbert, 2021; Depue et al., 2016). Based on these findings, it may be that the MTL - frontoparietal connectivity contributed to the suppression of competing information, which in turn led to the decreased sensitivity to similarity. Altogether, these results indicate an involvement of frontoparietal regions in resolving interference, however, more evidence is needed to elucidate whether and how these regions interact with the MTL to achieve interference resolution.

Although we chose to analyze the neural signals of pattern separation at the time of encoding in order to reduce the contaminating effects of retrieval, we cannot completely rule out the possibility that pattern separation and recognition processes took place during this phase. It is in fact likely that participants recognized repeat and similar fractals already during study, which may be also reflected in the activation of retrieval related areas, such as the angular gyrus, in the whole-brain results. However, participants were instructed to pay attention to the details of each trial (specifically fractal identity in the Object Task and fractal location in the Location Task) in order to commit them to their memory, and thus were encouraged to engage in the encoding, rather than retrieval of trials.

A limitation of this study is the combined analysis of the CA3 and dentate gyrus areas. Given the role of CA3 in pattern completion, one might argue that the activity pattern we see in the CA3DG is the result of pattern completion in the CA3. At the time of writing this work, there is no published valid and reliable method of delineating these two critical subfields on 3T MRI data (Olsen et al., 2019). In collapsing the two subfields, we have followed a substantial body of previous fMRI studies that nonetheless found a pattern separation signal in this combined region, albeit using everyday objects as stimuli (Chanales et al., 2017; Dimsdale-Zucker et al., 2018; Favila et al., 2016; Reagh & Yassa, 2014; Yassa & Stark, 2011).

To conclude, our work critically revisits the question of pattern separation in the CA3DG and domain-dependent interference reduction in the medial temporal neocortex. We replicated findings on content-specific processing in the MTL with non-meaningful stimuli, and provided evidence that frontoparietal regions contribute to mnemonic discrimination. Contrary to our expectations, we found no evidence that the CA3DG discriminates between repeats and interference-inducing similar stimuli. These results advance our understanding of how unique memories are created by a widespread collaboration of brain areas, and raise the intriguing question where the limits of hippocampal pattern separation lie.

## Supplementary Information

Supplementary information is available on OSF: https://osf.io/uzgx6

## Data and code availability

Neuroimaging data: https://openneuro.org/datasets/ds005559/ Behavioral data: https://osf.io/q9wks/?view_only=93849d7075374aad8dfce5b6c69e4e8a Analysis code: https://github.com/zs-nemecz/miniTRK

## Acknowledgements

This research was funded by the Hungarian National Research, Development and Innovation Office (FK128648). A.K. was additionally supported by a Max Planck Partner Group from the Max Planck Society, by a Bolyai János Research Scholarship of the Hungarian Academy of Sciences, and by the HUN-REN (grant 0708-21 515 AT). Parts of the work in this manuscript were conducted while Z.N. was a guest scientist at the Center for Lifespan Psychology, Max Planck Institute for Human Development, Berlin. Z.N. was supported by a One-Year Grant for Doctoral Candidates from the German Academic Exchange Service, and by the ÚNKP-22-3 New National Excellence Program of the Ministry for Culture and Innovation from the source of the National Research, Development and Innovation Fund. We are grateful for the services provided by the HUN-REN Cloud (Héder et al., 2022; https://science-cloud.hu/), where parts of the presented data were analyzed. We would like to thank Zoe Ngo, Myriam Sander, and Martin Dahl for helpful discussions about this project, and Eszter Somogyi and Dóra Bodócs for their assistance in data collection.

## References

Amer, T., & Davachi, L. (2023). Extra-hippocampal contributions to pattern separation. eLife, 12, e82250. 10.7554/eLife.82250

Anderson, M. C., & Hulbert, J. C. (2021). Active Forgetting: Adaptation of Memory by Prefrontal Control. Annual Review of Psychology, 72(Volume 72, 2021), 1–36. 10.1146/annurev-psych-072720-094140

Anderson, M. C., Ochsner, K. N., Kuhl, B., Cooper, J., Robertson, E., Gabrieli, S. W., Glover, G. H., & Gabrieli, J. D. E. (2004). Neural Systems Underlying the Suppression of Unwanted Memories. Science, 303(5655), 232–235. 10.1126/science.1089504

Avants, B. B., Epstein, C. L., Grossman, M., & Gee, J. C. (2008). Symmetric diffeomorphic image registration with cross-correlation: Evaluating automated labeling of elderly and neurodegenerative brain. Medical Image Analysis, 12(1), 26–41. 10.1016/j.media.2007.06.004

Avants, B. B., Tustison, N. J., Song, G., Cook, P. A., Klein, A., & Gee, J. C. (2011). A reproducible evaluation of ANTs similarity metric performance in brain image registration. NeuroImage, 54(3), 2033–2044. 10.1016/j.neuroimage.2010.09.025

Avants, B. B., Tustison, N., Song, G., & others. (2009). Advanced normalization tools (ANTS). Insight j, 2(365), 1–35.

Azab, M., Stark, S. M., & Stark, C. E. L. (2014). Contributions of human hippocampal subfields to spatial and temporal pattern separation. Hippocampus, 24(3), 293–302. 10.1002/hipo.22223

Baker, S., Vieweg, P., Gao, F., Gilboa, A., Wolbers, T., Black, S. E., & Rosenbaum, R. S. (2016). The Human Dentate Gyrus Plays a Necessary Role in Discriminating New Memories. Current Biology, 26(19), 2629–2634. 10.1016/j.cub.2016.07.081

Bakker, A., Kirwan, C. B., Miller, M., & Stark, C. E. L. (2008). Pattern Separation in the Human Hippocampal CA3 and Dentate Gyrus. Science, 319(5870), 1640–1642. 10.1126/science.1152882

Barense, M. D., Bussey, T. J., Lee, A. C. H., Rogers, T. T., Davies, R. R., Saksida, L. M., Murray, E. A., & Graham, K. S. (2005). Functional Specialization in the Human Medial Temporal Lobe. Journal of Neuroscience, 25(44), 10239–10246. 10.1523/JNEUROSCI.2704-05.2005

Barense, M. D., Henson, R. N. A., & Graham, K. S. (2011). Perception and Conception: Temporal Lobe Activity during Complex Discriminations of Familiar and Novel Faces and Objects. Journal of Cognitive Neuroscience, 23(10), 3052–3067. 10.1162/jocn_a_00010

Barense, M. D., Henson, R. N. A., Lee, A. C. H., & Graham, K. S. (2010). Medial temporal lobe activity during complex discrimination of faces, objects, and scenes: Effects of viewpoint. Hippocampus, 20(3), 389–401. 10.1002/hipo.20641

Bates, D., Mächler, M., Bolker, B., & Walker, S. (2015). Fitting Linear Mixed-Effects Models Using lme4. Journal of Statistical Software, 67(1), 1–48. 10.18637/jss.v067.i01

Bencze, D., Marián, M., Szőllősi, Á., Pajkossy, P., Nemecz, Z., Keresztes, A., Hermann, P., Vidnyánszky, Z., & Racsmány, M. (2024). Contribution of the lateral occipital and parahippocampal cortices to pattern separation of objects and contexts. Cerebral Cortex, 34(7), bhae295. 10.1093/cercor/bhae295

Bender, A. R., Keresztes, A., Bodammer, N. C., Shing, Y. L., Werkle-Bergner, M., Daugherty, A. M., Yu, Q., Kühn, S., Lindenberger, U., & Raz, N. (2018). Optimization and validation of automated hippocampal subfield segmentation across the lifespan. Human Brain Mapping, 39(2), 916–931. 10.1002/hbm.23891

Benjamini, Y., & Hochberg, Y. (1995). Controlling the False Discovery Rate: A Practical and Powerful Approach to Multiple Testing. Journal of the Royal Statistical Society: Series B (Methodological), 57(1), 289–300. 10.1111/j.2517-6161.1995.tb02031.x

Berron, D., Neumann, K., Maass, A., Schütze, H., Fliessbach, K., Kiven, V., Jessen, F., Sauvage, M., Kumaran, D., & Düzel, E. (2018). Age-related functional changes in domain-specific medial temporal lobe pathways. Neurobiology of Aging, 65, 86–97. 10.1016/j.neurobiolaging.2017.12.030

Berron, D., Schütze, H., Maass, A., Cardenas-Blanco, A., Kuijf, H. J., Kumaran, D., & Düzel, E. (2016). Strong Evidence for Pattern Separation in Human Dentate Gyrus. Journal of Neuroscience, 36(29), 7569–7579. 10.1523/JNEUROSCI.0518-16.2016

Brickman, A. M., Khan, U. A., Provenzano, F. A., Yeung, L.-K., Suzuki, W., Schroeter, H., Wall, M., Sloan, R. P., & Small, S. A. (2014). Enhancing dentate gyrus function with dietary flavanols improves cognition in older adults. Nature Neuroscience, 17(12), 1798–1803. 10.1038/nn.3850

Burke, S. N., Gaynor, L. S., Barnes, C. A., Bauer, R. M., Bizon, J. L., Roberson, E. D., & Ryan, L. (2018). Shared Functions of Perirhinal and Parahippocampal Cortices: Implications for Cognitive Aging. Trends in Neurosciences, 41(6), 349–359. 10.1016/j.tins.2018.03.001

Burke, S. N., Maurer, A. P., Nematollahi, S., Uprety, A., Wallace, J. L., & Barnes, C. A. (2014). Advanced Age Dissociates Dual Functions of the Perirhinal Cortex. Journal of Neuroscience, 34(2), 467–480. 10.1523/JNEUROSCI.2875-13.2014

Bussey, T. J., Saksida, L. M., & Murray, E. A. (2002). Perirhinal cortex resolves feature ambiguity in complex visual discriminations. European Journal of Neuroscience, 15(2), 365–374. 10.1046/j.0953-816x.2001.01851.x

Canada, K. L., Mazloum-Farzaghi, N., Rådman, G., Adams, J. N., Bakker, A., Baumeister, H., Berron, D., Bocchetta, M., Carr, V., Dalton, M. A., Flores, R. de, Keresztes, A., Joie, R. L., Mueller, S. G., Raz, N., Santini, T., Shaw, T., Stark, C. E. L., Tran, T. T., … Group, the H. S. (2023). A (Sub)field Guide to Quality Control in Hippocampal Subfield Segmentation on High-resolution T2-weighted MRI (p. 2023.11.29.568895). bioRxiv. 10.1101/2023.11.29.568895

Cayco-Gajic, N. A., & Silver, R. A. (2019). Re-evaluating Circuit Mechanisms Underlying Pattern Separation. Neuron, 101(4), 584–602. 10.1016/j.neuron.2019.01.044

Chanales, A. J. H., Oza, A., Favila, S. E., & Kuhl, B. A. (2017). Overlap among Spatial Memories Triggers Repulsion of Hippocampal Representations. Current Biology, 27(15), 2307–2317.e5. 10.1016/j.cub.2017.06.057

Connor, C. E., & Knierim, J. J. (2017). Integration of objects and space in perception and memory. Nature Neuroscience, 20(11), 1493–1503. 10.1038/nn.4657

Cowell, R. A., Barense, M. D., & Sadil, P. S. (2019). A Roadmap for Understanding Memory: Decomposing Cognitive Processes into Operations and Representations. eNeuro, 6(4). 10.1523/ENEURO.0122-19.2019

Dale, A., Fischl, B., & Sereno, M. I. (1999). Cortical Surface-Based Analysis: I. Segmentation and Surface Reconstruction. NeuroImage, 9(2), 179–194.

Depue, B. E., Orr, J. M., Smolker, H. R., Naaz, F., & Banich, M. T. (2016). The Organization of Right Prefrontal Networks Reveals Common Mechanisms of Inhibitory Regulation Across Cognitive, Emotional, and Motor Processes. Cerebral Cortex, 26(4), 1634–1646. 10.1093/cercor/bhu324

Desikan, R. S., Ségonne, F., Fischl, B., Quinn, B. T., Dickerson, B. C., Blacker, D., Buckner, R. L., Dale, A. M., Maguire, R. P., Hyman, B. T., Albert, M. S., & Killiany, R. J. (2006). An automated labeling system for subdividing the human cerebral cortex on MRI scans into gyral based regions of interest. NeuroImage, 31(3), 968–980. 10.1016/j.neuroimage.2006.01.021

Destrieux, C., Fischl, B., Dale, A., & Halgren, E. (2010). Automatic parcellation of human cortical gyri and sulci using standard anatomical nomenclature. NeuroImage, 53(1), 1–15. 10.1016/j.neuroimage.2010.06.010

Di, X., & Biswal, B. B. (2017). Psychophysiological Interactions in a Visual Checkerboard Task: Reproducibility, Reliability, and the Effects of Deconvolution. Frontiers in Neuroscience, 11. 10.3389/fnins.2017.00573

Di, X., Zhang, Z., & Biswal, B. B. (2021). Understanding psychophysiological interaction and its relations to beta series correlation. Brain Imaging and Behavior, 15(2), 958–973. 10.1007/s11682-020-00304-8

Dimsdale-Zucker, H. R., Ritchey, M., Ekstrom, A. D., Yonelinas, A. P., & Ranganath, C. (2018). CA1 and CA3 differentially support spontaneous retrieval of episodic contexts within human hippocampal subfields. Nature Communications, 9(1), Article 1. 10.1038/s41467-017-02752-1

Dixon, M. L., De La Vega, A., Mills, C., Andrews-Hanna, J., Spreng, R. N., Cole, M. W., & Christoff, K. (2018). Heterogeneity within the frontoparietal control network and its relationship to the default and dorsal attention networks. Proceedings of the National Academy of Sciences, 115(7), E1598–E1607. 10.1073/pnas.1715766115

Duff, M. C., Warren, D. E., Gupta, R., Vidal, J. P. B., Tranel, D., & Cohen, N. J. (2012). Teasing apart tangrams: Testing hippocampal pattern separation with a collaborative referencing paradigm. Hippocampus, 22(5), 1087–1091. 10.1002/hipo.20967

Eacott, M. J., & Gaffan, E. A. (2005). The Roles of Perirhinal Cortex, Postrhinal Cortex, and the Fornix in Memory for Objects, Contexts, and Events in the Rat. The Quarterly Journal of Experimental Psychology Section B, 58(3–4b), 202–217. 10.1080/02724990444000203

Favila, S. E., Chanales, A. J. H., & Kuhl, B. A. (2016). Experience-dependent hippocampal pattern differentiation prevents interference during subsequent learning. Nature Communications, 7(1), 11066. 10.1038/ncomms11066

Fischl, B. (2004). Automatically Parcellating the Human Cerebral Cortex. Cerebral Cortex, 14(1), 11–22. 10.1093/cercor/bhg087

Fischl, B., Salat, D. H., Busa, E., Albert, M., Dieterich, M., Haselgrove, C., van der Kouwe, A., Killiany, R., Kennedy, D., Klaveness, S., Montillo, A., Makris, N., Rosen, B., & Dale, A. M. (2002). Whole brain segmentation: Automated labeling of neuroanatomical structures in the human brain. Neuron, 33, 341–355.

Fischl, B., Sereno, M. I., & Dale, A. (1999). Cortical Surface-Based Analysis: II: Inflation, Flattening, and a Surface-Based Coordinate System. NeuroImage, 9(2), 195–207.

Frazier, J. A., Chiu, S., Breeze, J. L., Makris, N., Lange, N., Kennedy, D. N., Herbert, M. R., Bent, E. K., Koneru, V. K., Dieterich, M. E., Hodge, S. M., Rauch, S. L., Grant, P. E., Cohen, B. M., Seidman, L. J., Caviness, V. S., & Biederman, J. (2005). Structural Brain Magnetic Resonance Imaging of Limbic and Thalamic Volumes in Pediatric Bipolar Disorder. American Journal of Psychiatry, 162(7), 1256–1265. 10.1176/appi.ajp.162.7.1256

Friston, K. J., Buechel, C., Fink, G. R., Morris, J., Rolls, E., & Dolan, R. J. (1997). Psychophysiological and Modulatory Interactions in Neuroimaging. NeuroImage, 6(3), 218–229. 10.1006/nimg.1997.0291

Gilbert, P. E., Kesner, R. P., & Lee, I. (2001). Dissociating hippocampal subregions: A double dissociation between dentate gyrus and CA1. Hippocampus, 11(6), 626–636. 10.1002/hipo.1077

Goldstein, J. M., Seidman, L. J., Makris, N., Ahern, T., O’Brien, L. M., Caviness, V. S., Kennedy, D. N., Faraone, S. V., & Tsuang, M. T. (2007). Hypothalamic Abnormalities in Schizophrenia: Sex Effects and Genetic Vulnerability. Biological Psychiatry, 61(8), 935–945. 10.1016/j.biopsych.2006.06.027

GoodSmith, D., Lee, H., Neunuebel, J. P., Song, H., & Knierim, J. J. (2019). Dentate Gyrus Mossy Cells Share a Role in Pattern Separation with Dentate Granule Cells and Proximal CA3 Pyramidal Cells. Journal of Neuroscience, 39(48), 9570–9584. 10.1523/JNEUROSCI.0940-19.2019

Gorgolewski, K. J., Varoquaux, G., Rivera, G., Schwartz, Y., Sochat, V. V., Ghosh, S. S., Maumet, C., Nichols, T. E., Poline, J.-B., Yarkoni, T., Margulies, D. S., & Poldrack, R. A. (2016). NeuroVault.org: A repository for sharing unthresholded statistical maps, parcellations, and atlases of the human brain. NeuroImage, 124(Pt B), 1242–1244. 10.1016/j.neuroimage.2015.04.016

Greve, D. N., & Fischl, B. (2009). Accurate and robust brain image alignment using boundary-based registration. NeuroImage, 48(1), 63–72. 10.1016/j.neuroimage.2009.06.060

Hunsaker, M. R., Rosenberg, J. S., & Kesner, R. P. (2008). The role of the dentate gyrus, CA3a,b, and CA3c for detecting spatial and environmental novelty. Hippocampus, 18(10), 1064–1073. 10.1002/hipo.20464

Hunter, J. D. (2007). Matplotlib: A 2D graphics environment. Computing in Science & Engineering, 9(3), 90–95. 10.1109/MCSE.2007.55

Jenkinson, M., Bannister, P., Brady, M., & Smith, S. (2002). Improved Optimization for the Robust and Accurate Linear Registration and Motion Correction of Brain Images. NeuroImage, 17(2), 825–841. 10.1006/nimg.2002.1132

Jenkinson, M., Beckmann, C. F., Behrens, T. E. J., Woolrich, M. W., & Smith, S. M. (2012). FSL. NeuroImage, 62(2), 782–790. 10.1016/j.neuroimage.2011.09.015

Jenkinson, M., & Smith, S. (2001). A global optimisation method for robust affine registration of brain images. Medical Image Analysis, 5(2), 143–156. 10.1016/S1361-8415(01)00036-6

Kirwan, C. B., & Stark, C. E. L. (2007). Overcoming interference: An fMRI investigation of pattern separation in the medial temporal lobe. Learning & Memory, 14(9), 625–633. 10.1101/lm.663507

Klippenstein, J. L., Stark, S. M., Stark, C. E. L., & Bennett, I. J. (2020). Neural substrates of mnemonic discrimination: A whole-brain fMRI investigation. Brain and Behavior, 10(3), e01560. 10.1002/brb3.1560

Knierim, J. J., & Neunuebel, J. P. (2016). Tracking the flow of hippocampal computation: Pattern separation, pattern completion, and attractor dynamics. Neurobiology of Learning and Memory, 129, 38–49. 10.1016/j.nlm.2015.10.008

Kyle, C. T., Stokes, J. D., Lieberman, J. S., Hassan, A. S., & Ekstrom, A. D. (2015). Successful retrieval of competing spatial environments in humans involves hippocampal pattern separation mechanisms. eLife, 4, e10499. 10.7554/eLife.10499

Lawrence, A. V., Cardoza, J., & Ryan, L. (2020). Medial temporal lobe regions mediate complex visual discriminations for both objects and scenes: A process-based view. Hippocampus, hipo.23203. 10.1002/hipo.23203

Leal, S. L., & Yassa, M. A. (2018). Integrating new findings and examining clinical applications of pattern separation. Nature Neuroscience, 21(2), Article 2. 10.1038/s41593-017-0065-1

Leutgeb, J. K., Leutgeb, S., Moser, M.-B., & Moser, E. I. (2007). Pattern Separation in the Dentate Gyrus and CA3 of the Hippocampus. Science, 315(5814), 961–966. 10.1126/science.1135801

Liu, K. Y., Gould, R. L., Coulson, M. C., Ward, E. V., & Howard, R. J. (2016). Tests of pattern separation and pattern completion in humans—A systematic review. Hippocampus, 26(6), 705–717. 10.1002/hipo.22561

Loiotile, R. E., & Courtney, S. M. (2015). A signal detection theory analysis of behavioral pattern separation paradigms. Learning & Memory, 22(8), 364–369. 10.1101/lm.038141.115

Makris, N., Goldstein, J. M., Kennedy, D., Hodge, S. M., Caviness, V. S., Faraone, S. V., Tsuang, M. T., & Seidman, L. J. (2006). Decreased volume of left and total anterior insular lobule in schizophrenia. Schizophrenia Research, 83(2), 155–171. 10.1016/j.schres.2005.11.020

Martin, C. B., & Barense, M. D. (2023). Perception and Memory in the Ventral Visual Stream and Medial Temporal Lobe. Annual Review of Vision Science, 9(1), 409–434. 10.1146/annurev-vision-120222-014200

McHugh, T. J., Jones, M. W., Quinn, J. J., Balthasar, N., Coppari, R., Elmquist, J. K., Lowell, B. B., Fanselow, M. S., Wilson, M. A., & Tonegawa, S. (2007). Dentate Gyrus NMDA Receptors Mediate Rapid Pattern Separation in the Hippocampal Network. Science, 317(5834), 94–99. 10.1126/science.1140263

McLaren, D. G., Ries, M. L., Xu, G., & Johnson, S. C. (2012). A Generalized Form of Context-Dependent Psychophysiological Interactions (gPPI): A Comparison to Standard Approaches. Neuroimage, 61(4), 1277–1286. 10.1016/j.neuroimage.2012.03.068

Mitchnick, K. A., Ahmad, Z., Mitchnick, S. D., Ryan, J. D., Rosenbaum, R. S., & Freud, E. (2022). Damage to the human dentate gyrus impairs the perceptual discrimination of complex, novel objects. Neuropsychologia, 172, 108238. 10.1016/j.neuropsychologia.2022.108238

Mitchnick, K. A., Marlatte, H., Belchev, Z., Gao, F., & Rosenbaum, R. S. (2024). Differential contributions of the hippocampal dentate gyrus and CA1 subfield to mnemonic discrimination. Hippocampus, 34(6), 278–283. 10.1002/hipo.23604

Motley, S. E., & Kirwan, C. B. (2012). A Parametric Investigation of Pattern Separation Processes in the Medial Temporal Lobe. Journal of Neuroscience, 32(38), 13076–13084. 10.1523/JNEUROSCI.5920-11.2012

Mumford, J. A., Bissett, P. G., Jones, H. M., Shim, S., Rios, J. A. H., & Poldrack, R. A. (2024). The response time paradox in functional magnetic resonance imaging analyses. Nature Human Behaviour, 8(2), 349–360. 10.1038/s41562-023-01760-0

Nash, M. I., Hodges, C. B., Muncy, N. M., & Kirwan, C. B. (2021). Pattern separation beyond the hippocampus: A high-resolution whole-brain investigation of mnemonic discrimination in healthy adults. Hippocampus, 31(4), 408–421. 10.1002/hipo.23299

Neunuebel, J. P., & Knierim, J. J. (2014). CA3 retrieves coherent representations from degraded input: Direct evidence for CA3 pattern completion and dentate gyrus pattern separation. Neuron, 81(2), 416–427. 10.1016/j.neuron.2013.11.017

Nilssen, E. S., Doan, T. P., Nigro, M. J., Ohara, S., & Witter, M. P. (2019). Neurons and networks in the entorhinal cortex: A reappraisal of the lateral and medial entorhinal subdivisions mediating parallel cortical pathways. Hippocampus, 29(12), 1238–1254. 10.1002/hipo.23145

Olsen, R. K., Carr, V. A., Daugherty, A. M., La Joie, R., Amaral, R. S. C., Amunts, K., Augustinack, J. C., Bakker, A., Bender, A. R., Berron, D., Boccardi, M., Bocchetta, M., Burggren, A. C., Chakravarty, M. M., Chételat, G., de Flores, R., DeKraker, J., Ding, S.-L., Geerlings, M. I., … Group, H. S. (2019). Progress update from the hippocampal subfields group. Alzheimer’s & Dementia: Diagnosis, Assessment & Disease Monitoring, 11(1), 439–449. 10.1016/j.dadm.2019.04.001

O’Reilly, J. X., Woolrich, M. W., Behrens, T. E. J., Smith, S. M., & Johansen-Berg, H. (2012). Tools of the trade: Psychophysiological interactions and functional connectivity. Social Cognitive and Affective Neuroscience, 7(5), 604–609. 10.1093/scan/nss055

Paleja, M., Girard, T. A., Herdman, K. A., & Christensen, B. K. (2014). Two distinct neural networks functionally connected to the human hippocampus during pattern separation tasks. Brain and Cognition, 92, 101–111. 10.1016/j.bandc.2014.10.009

Peirce, J., Gray, J. R., Simpson, S., MacAskill, M., Höchenberger, R., Sogo, H., Kastman, E., & Lindeløv, J. K. (2019). PsychoPy2: Experiments in behavior made easy. Behavior Research Methods, 51(1), 195–203. 10.3758/s13428-018-01193-y

Quiroga, R. Q. (2020). No Pattern Separation in the Human Hippocampus. Trends in Cognitive Sciences, 24(12), 994–1007. 10.1016/j.tics.2020.09.012

Reagh, Z. M., & Yassa, M. A. (2014). Object and spatial mnemonic interference differentially engage lateral and medial entorhinal cortex in humans. Proceedings of the National Academy of Sciences, 111(40), E4264–E4273. 10.1073/pnas.1411250111

Rolls, E. T. (2013). The mechanisms for pattern completion and pattern separation in the hippocampus. Frontiers in Systems Neuroscience, 7. 10.3389/fnsys.2013.00074

Rolls, E. T. (2024). Two what, two where, visual cortical streams in humans. Neuroscience & Biobehavioral Reviews, 160, 105650. 10.1016/j.neubiorev.2024.105650

Seabold, S., & Perktold, J. (2010). statsmodels: Econometric and statistical modeling with python. 9th Python in Science Conference.

Staresina, B. P., Duncan, K. D., & Davachi, L. (2011). Perirhinal and Parahippocampal Cortices Differentially Contribute to Later Recollection of Object- and Scene-Related Event Details. Journal of Neuroscience, 31(24), 8739–8747. 10.1523/JNEUROSCI.4978-10.2011

Stark, S. M., Stevenson, R., Wu, C., Rutledge, S., & Stark, C. E. L. (2015). Stability of age-related deficits in the mnemonic similarity task across task variations. Behavioral Neuroscience, 129(3), 257–268. 10.1037/bne0000055

Stark, S. M., Yassa, M. A., Lacy, J. W., & Stark, C. E. L. (2013). A task to assess behavioral pattern separation (BPS) in humans: Data from healthy aging and mild cognitive impairment. Neuropsychologia, 51(12), 2442–2449. 10.1016/j.neuropsychologia.2012.12.014

Uddin, L. Q., Yeo, B. T. T., & Spreng, R. N. (2019). Towards a universal taxonomy of macro-scale functional human brain networks. Brain Topography, 32(6), 926. 10.1007/s10548-019-00744-6

Virtanen, P., Gommers, R., Oliphant, T. E., Haberland, M., Reddy, T., Cournapeau, D., Burovski, E., Peterson, P., Weckesser, W., Bright, J., van der Walt, S. J., Brett, M., Wilson, J., Millman, K. J., Mayorov, N., Nelson, A. R. J., Jones, E., Kern, R., Larson, E., … SciPy 1.0 Contributors. (2020). SciPy 1.0: Fundamental Algorithms for Scientific Computing in Python. Nature Methods, 17, 261–272. 10.1038/s41592-019-0686-2

Wammes, J. D., Norman, K. A., & Turk-Browne, N. B. (2021). Increasing stimulus similarity drives nonmonotonic representational change in hippocampus. bioRxiv, 2021.03.13.435275. 10.1101/2021.03.13.435275

Wanjia, G., Favila, S. E., Kim, G., Molitor, R. J., & Kuhl, B. A. (2021). Abrupt hippocampal remapping signals resolution of memory interference. Nature Communications, 12(1), 4816. 10.1038/s41467-021-25126-0

Waskom, M. L. (2021). seaborn: Statistical data visualization. Journal of Open Source Software, 6(60), 3021. 10.21105/joss.03021

Wimber, M., Alink, A., Charest, I., Kriegeskorte, N., & Anderson, M. C. (2015). Retrieval induces adaptive forgetting of competing memories via cortical pattern suppression. Nature Neuroscience, 18(4), 582–589. 10.1038/nn.3973

Wisse, L. E. M., Daugherty, A. M., Olsen, R. K., Berron, D., Carr, V. A., Stark, C. E. L., Amaral, R. S. C., Amunts, K., Augustinack, J. C., Bender, A. R., Bernstein, J. D., Boccardi, M., Bocchetta, M., Burggren, A., Chakravarty, M. M., Chupin, M., Ekstrom, A., de Flores, R., Insausti, R., … Group, for the H. S. (2017). A harmonized segmentation protocol for hippocampal and parahippocampal subregions: Why do we need one and what are the key goals? Hippocampus, 27(1), 3–11. 10.1002/hipo.22671

Woolrich, M. W., Behrens, T. E. J., Beckmann, C. F., Jenkinson, M., & Smith, S. M. (2004). Multilevel linear modelling for FMRI group analysis using Bayesian inference. NeuroImage, 21(4), 1732–1747. 10.1016/j.neuroimage.2003.12.023

Woolrich, M. W., Jbabdi, S., Patenaude, B., Chappell, M., Makni, S., Behrens, T., Beckmann, C., Jenkinson, M., & Smith, S. M. (2009). Bayesian analysis of neuroimaging data in FSL. NeuroImage, 45(1, Supplement 1), S173–S186. 10.1016/j.neuroimage.2008.10.055

Woolrich, M. W., Ripley, B. D., Brady, M., & Smith, S. M. (2001). Temporal Autocorrelation in Univariate Linear Modeling of FMRI Data. NeuroImage, 14(6), 1370–1386. 10.1006/nimg.2001.0931

Yarkoni, T., Poldrack, R. A., Nichols, T. E., Van Essen, D. C., & Wager, T. D. (2011). Large-scale automated synthesis of human functional neuroimaging data. Nature Methods, 8(8), 665–670. 10.1038/nmeth.1635

Yassa, M. A., Mattfeld, A. T., Stark, S. M., & Stark, C. E. L. (2011). Age-related memory deficits linked to circuit-specific disruptions in the hippocampus. Proceedings of the National Academy of Sciences, 108(21), 8873–8878. 10.1073/pnas.1101567108

Yassa, M. A., & Stark, C. E. L. (2011). Pattern separation in the hippocampus. Trends in Neurosciences, 34(10), 515–525. 10.1016/j.tins.2011.06.006

Yonelinas, A. P. (2013). The hippocampus supports high-resolution binding in the service of perception, working memory and long-term memory. Behavioural Brain Research, 254, 34–44. 10.1016/j.bbr.2013.05.030

Yushkevich, P. A., Pluta, J. B., Wang, H., Xie, L., Ding, S.-L., Gertje, E. C., Mancuso, L., Kliot, D., Das, S. R., & Wolk, D. A. (2015). Automated volumetry and regional thickness analysis of hippocampal subfields and medial temporal cortical structures in mild cognitive impairment. Human Brain Mapping, 36(1), 258–287. 10.1002/hbm.22627

