## Supplementary Material for "fMRI signals of pattern separation in the neocortex and hippocampus to non-meaningful objects and their spatial location"

Supplementary Figure 7 ... 13
Object Similarity Rating Experiment ... 13
Supplementary Figure 8 ... 14
Segmentation of region of interests in the medial temporal lobe ... 14
Supplementary Figure 9. Medial temporal ROIs on an example segmentations. 14
Registration of partial FOV high-resolution PD images to functional scans ... 15
Supplementary Figure 10. Example ROI to Functional registrations ... 16
Mixed effects model assumptions ... 17
Behavioral Models: Generalized binomial mixed effects models ... 17
Neural Models: Linear mixed effects models ... 17
Supplementary Figure 11 ... 17
References ... 18

### Supplementary Results

#### Models of behavioral sensitivity to the first and second object similarity

**Supplementary Table 1. The effect of similarity on the ratio of "new" responses in the Object Task.**

| Model | Fixed effects | Estimate | Std Error | *z* | *p* | AIC |
|---|---|---|---|---|---|---|
| **A** | *FirstSimilarity* | 0.16 | 0.02 | 9.23 | < .001 | 2562.93 |
| **B** | *SecondSimilarity* | 0.17 | 0.02 | 8.98 | < .001 | 2565.45 |
| **C** | *FirstSimilarity* | 0.02 | 0.06 | 0.3 | .76 | 2545.91 |
| | *SecondSimilarity* | 0.09 | 0.03 | 3.41 | < .001 | |
| | *FirstS. * SecondS.* | 0.02 | 0.01 | 1.64 | .1 | |

**Supplementary Table 2.** Model Comparisons - The effect of object feature similarity on the ratio of new responses in the Object Task

| Models | N par. | AIC | BIC | Log Likelihood | Deviance | Chi-square | df | *p* |
|---|---|---|---|---|---|---|---|---|
| **A** | 3 | 2562.9 | 2579.8 | -1278.5 | 2556.9 | | 2 | < .001 |
| **C** | 5 | 2545.9 | 2574.1 | -1268.0 | 2535.9 | 21.02 | | |
| **B** | 3 | 2565.5 | 2582.4 | -1279.7 | 2559.5 | | | < .001 |
| **C** | 5 | 2545.9 | 2574.1 | -1268.0 | 2535.9 | 23.55 | 2 | |

#### Models of behavioral sensitivity to the first and second spatial similarity

**Supplementary Table 3. The effect of similarity on the ratio of "new" responses in the Location Task.**

| **Model** | **Fixed effects** | **Estimate** | **Std Error** | ***z*** | ***p*** | **AIC** |
|---|---|---|---|---|---|---|
| **A** | *FirstSimilarity* | 0.11 | 0.02 | 7 | < .001 | 4242.24 |
| **B** | *SecondSimilarity* | 0.12 | 0.02 | 6.5 | < .001 | 4249.73 |
| | *FirstSimilarity* | 0.23 | 0.05 | 4.53 | < .001 | |
| **C** | *SecondSimilarity* | 0.11 | 0.04 | 2.74 | .006 | 4233.07 |
| | *FirstS. * SecondS.* | -0.03 | 0.01 | -3.49 | < .001 | |

**Supplementary Table 4.** Model Comparisons - The effect of spatial similarity on the ratio of new responses in the Location Task

| Models | N par. | AIC | BIC | Log Likelihood | Deviance | Chi-square | df | *p* |
|---|---|---|---|---|---|---|---|---|
| **A** | 3 | 4242.2 | 4260.4 | -2118.1 | 4236.2 | | | |
| **C** | 5 | 4233.1 | 4263.3 | -2111.5 | 4223.1 | 13.16 | 2 | .001 |
| **B** | 3 | 4249.7 | 4267.8 | -2121.9 | 4243.7 | | | |
| **C** | 5 | 4233.1 | 4263.3 | -2111.5 | 4223.2 | 20.67 | 2 | < .001 |

#### Associations between behavior and neuronal sensitivity to similarity

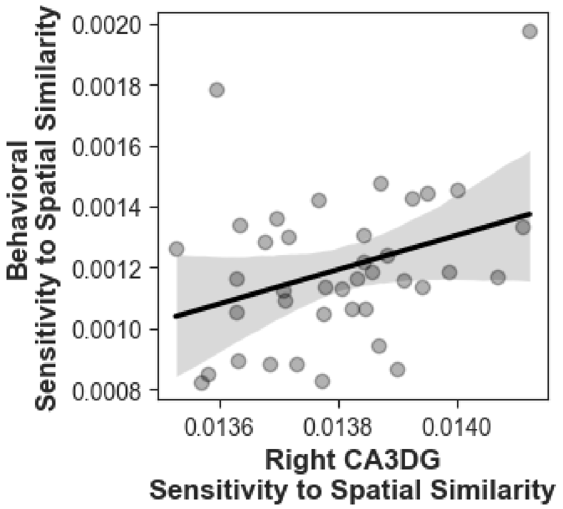

**Supplementary Figure 1. The association between the right CA3DG sensitivity to spatial similarity and behavioral sensitivity to spatial sensitivity**

#### Modeling Baseline (multiple repetition) trials

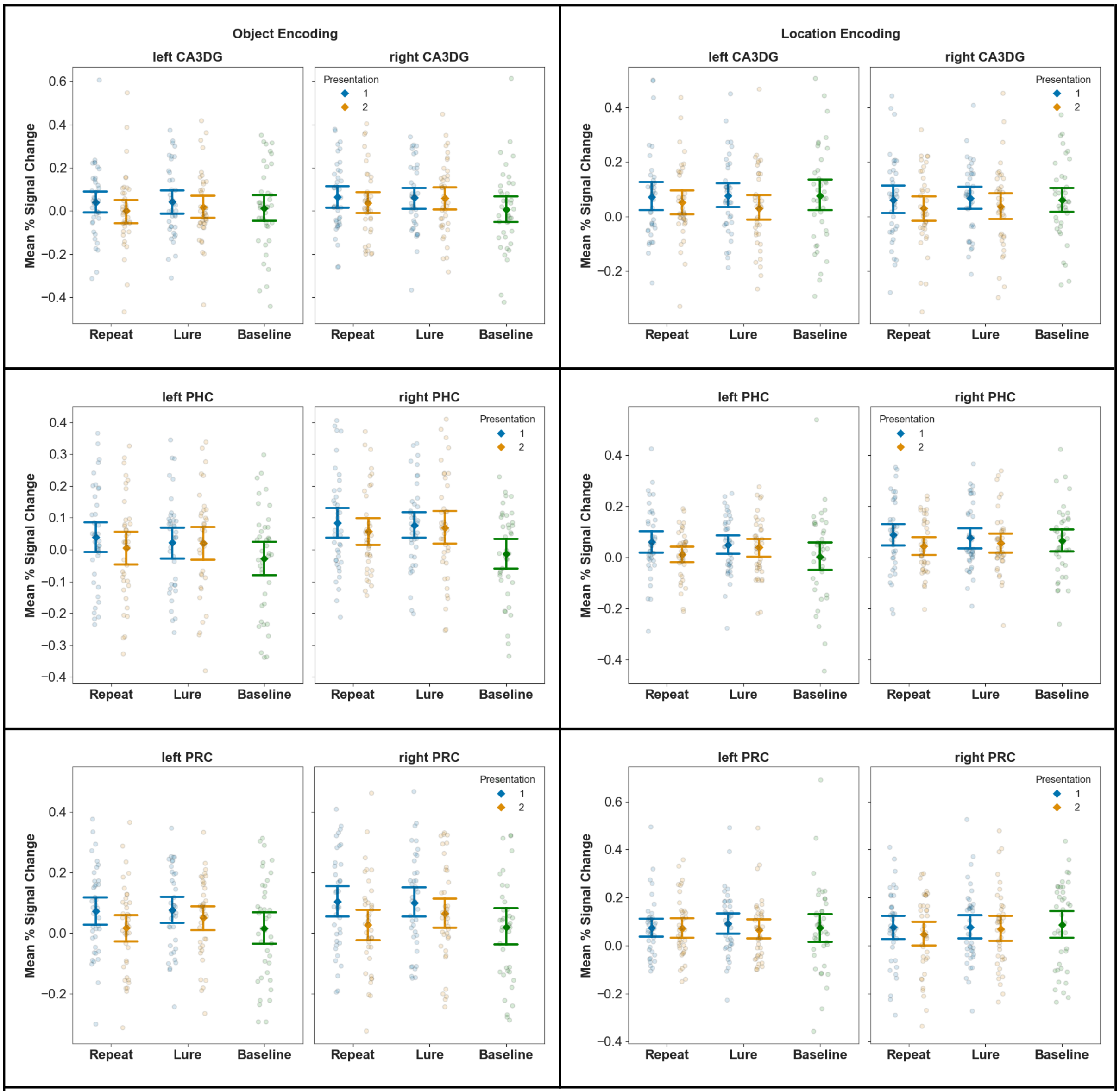

**Supplementary Figure 2.**

#### Whole-brain analysis: (*Similar - First*) > (*Repeats - First*) contrast results

**Supplementary Table 4.** Significant clusters for contrast (*Similar - First*) > (*Repeats - First*) during Object Encoding

| Area | N voxels | Max. Z value | *p* |
|---|---|---|---|
| bilateral lateral occipital cortex, occipital fusiform gyrus, right inferior temporal gyrus | 25676 | 4.97 | < .001 |
| right inferior frontal gyrus, right precentral gyrus, right middle frontal gyrus | 1609 | 3.79 | < .001 |
| left lateral occipital cortex | 1456 | 3.52 | < .001 |
| right lateral occipital cortex | 1349 | 3.63 | < .001 |
| right postcentral gyrus and right supramarginal gyrus | 980 | 3.27 | < .001 |
| left frontal pole, left inferior frontal gyrus (pars triangularis) | 891 | 3.45 | < .001 |
| bilateral paracingulate cortex | 821 | 3.51 | < .001 |
| left inferior frontal gyrus (pars opercularis) | 723 | 3.85 | < .001 |
| right postcentral gyrus, and right superior parietal lobule | 596 | 3.26 | < .001 |
| right frontal operculum and insular cortex, right putamen | 595 | 3.23 | < .001 |
| left postcentral gyrus and left supramarginal gyrus | 542 | 3.33 | < .001 |
| left precentral gyrus and middle frontal gyrus (frontal eye field) | 488 | 3.49 | < .001 |
| right frontal pole | 473 | 3.87 | < .001 |
| left postcentral gyrus and left supramarginal gyrus | 449 | 3.56 | < .001 |
| left superior parietal lobule | 405 | 3.37 | .002 |
| bilateral supplementary motor area | 289 | 3.4 | .034 |

**Supplementary Table 5.** Significant clusters for contrast (*Similar - First*) > (*Repeats - First*) during Location Encoding

| Area | N voxels | Max. Z value | *p* |
|---|---|---|---|
| bilateral lingual gyrus, bilateral occipital fusiform gyrus, bilateral occipital pole and lateral occipital cortex | 8124 | 4.46 | < .001 |

#### Whole-brain analysis: *Similar - First* contrast results

**Supplementary Table 6.** Significant clusters for the contrast *Similar - First* during Object Encoding

| Area | N voxels | Max. Z value | *p* |
|---|---|---|---|
| right frontal pole, right middle frontal gyrus, right orbitofrontal cortex, bilateral paracingulate gyrus | 31370 | 5.91 | < .001 |
| bilateral precuneus | 15531 | 5.63 | < .001 |
| right lateral occipital cortex, right angular gyrus, right supramarginal gyrus | 13958 | 6.2 | < .001 |
| left lateral occipital cortex, left supramarginal gyrus, left angular gyrus | 8268 | 5.18 | < .001 |
| left middle frontal gyrus | 6515 | 4.87 | < .001 |
| left frontal pole | 4505 | 5.18 | < .001 |
| right middle temporal gyrus (posterior division) | 4051 | 4.38 | < .001 |
| left middle temporal gyrus, posterior division | 1552 | 4.26 | < .001 |
| bilateral occipital pole and lingual gyrus | 1466 | 3.46 | < .001 |
| bilateral posterior cingulate | 970 | 4.34 | < .001 |
| left cerebellum | 665 | 5.41 | < .001 |
| left orbitofrontal cortex | 647 | 4.27 | < .001 |
| right cerebellum | 301 | 3.55 | .034 |

**Supplementary Table 7.** Significant clusters for the contrast *Similar - First* during Location Encoding

| Area | N voxels | Max. Z value | *p* |
|---|---|---|---|
| right angular gyrus, right supramarginal gyrus, bilateral precuneus, lingual gyrus, occipital pole, bilateral lateral occipital cortex | 44544 | 5.68 | < .001 |
| right middle frontal gyrus | 9691 | 4.71 | < .001 |
| left angular gyrus, supramarginal gyrus, left lateral occipital cortex | 5509 | 4.26 | < .001 |
| left middle frontal gyrus | 5457 | 4.28 | < .001 |
| right frontal pole | 5156 | 4.6 | < .001 |
| left frontal pole | 3657 | 4.44 | < .001 |
| right superior and middle temporal gyrus | 2316 | 4.19 | < .001 |
| bilateral superior frontal gyrus and paracingulate gyrus | 1924 | 04.04 | < .001 |
| left superior, middle, and inferior temporal gyrus | 944 | 3.67 | < .001 |
| left cerebellum | 553 | 4 | < .001 |
| right frontal pole | 456 | 3.48 | < .001 |
| right orbitofrontal cortex | 360 | 3.57 | .007 |

#### Neurosynth Analysis

| Supplementary Table 8. Object Similar > First. Top 10 Results. | |
|---|---|
| **Neurosynth term** | **Correlation** |
| parietal | 0.199 |
| inferior parietal | 0.189 |
| dorsolateral | 0.183 |
| working memory | 0.166 |
| working | 0.164 |
| parietal | 0.164 |
| precuneus | 0.158 |
| dorsolateral prefrontal | 0.146 |
| frontoparietal | 0.143 |
| task | 0.138 |

| Supplementary Table 9. Object Similar > First. Memory related terms. | |
|---|---|
| **Neurosynth term** | **Correlation** |
| default mode | 0.026 |
| memory retrieval | 0.104 |
| recognition | 0.07 |
| episodic memory | -0.001 |

| Supplementary Table 10. Location Similar > First. Top 10 Results. | |
|---|---|
| **Neurosynth term** | **Correlation** |
| parietal | 0.232 |
| precuneus | 0.189 |
| parietal cortex | 0.188 |
| working memory | 0.183 |
| working | 0.181 |

| | |
|---|---|
| inferior parietal | 0.167 |
| cuneus | 0.158 |
| frontoparietal | 0.154 |
| dorsolateral | 0.138 |
| memory | 0.136 |

| **Supplementary Table 11. Location Similar > First. Memory related terms.** | |
|---|---|
| **Neurosynth term** | **Correlation** |
| default mode | 0.042 |
| memory retrieval | 0.09 |
| recognition | 0.06 |
| episodic memory | 0.014 |

#### Connectivity between the frontoparietal and medial temporal regions

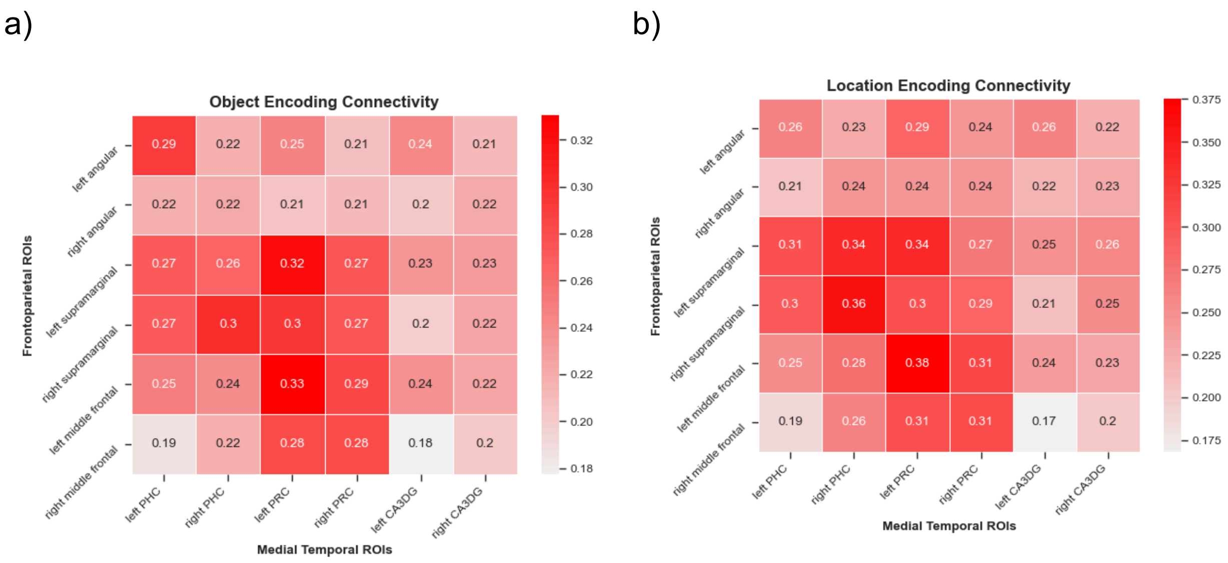

**Supplementary Figure 3.**

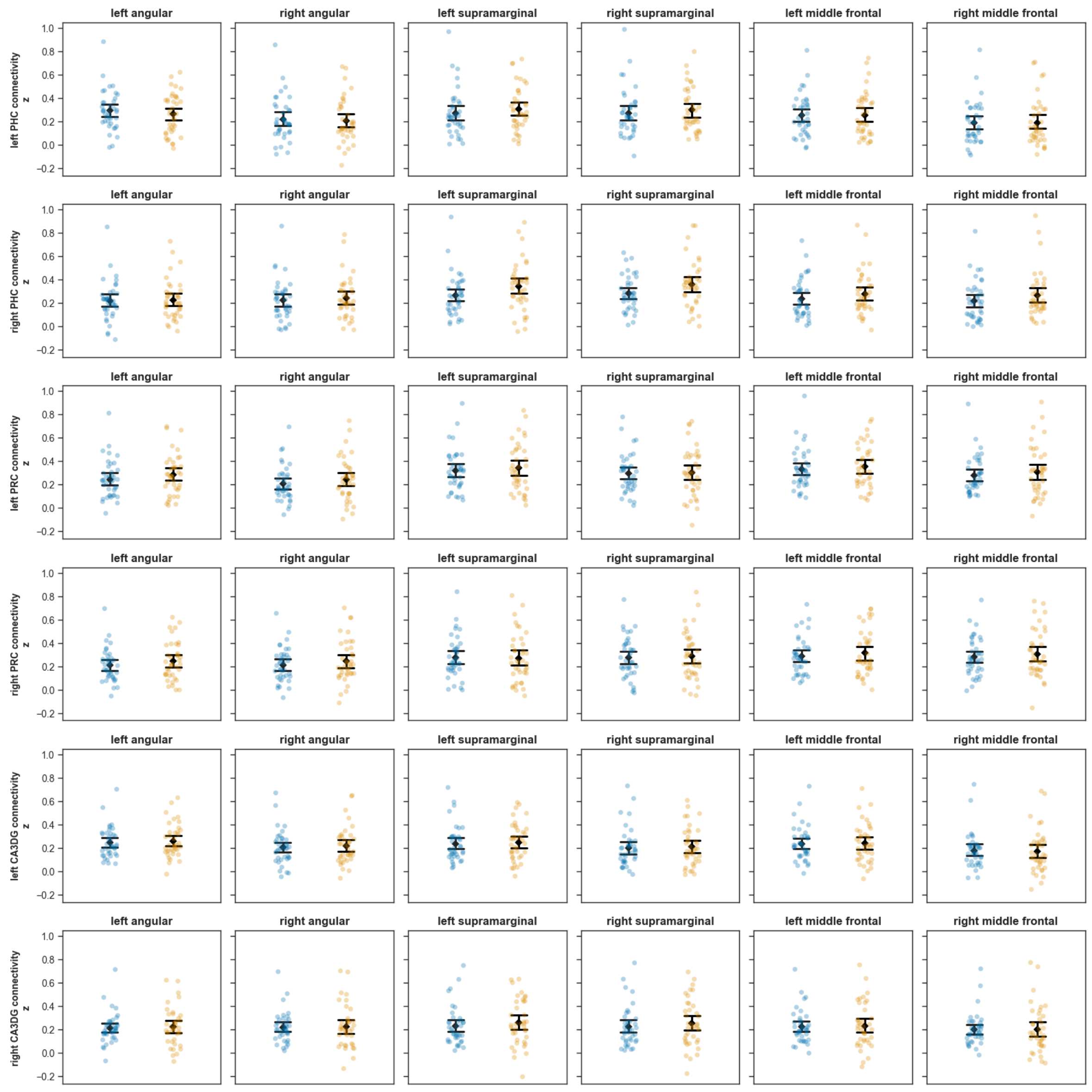

**Supplementary Figure 4.**

#### Associations between behavior and frontoparietal-medial temporal connectivity

**Supplementary Table 12. Object mnemonic discrimination and frontoparietal-medial temporal connectivity**

| Similarity Sensitivity | Task | MTL ROI | Frontoparietal ROI | Spearman r | p |
|---|---|---|---|---|---|
| SecondSimilarity | Object | left PRC | right supramarginal | -0.41 | 0.001 |
| SecondSimilarity | Object | right PHC | left supramarginal | -0.41 | 0.001 |
| FirstSimilarity | Object | left PRC | right angular | -0.41 | 0.001 |
| FirstSimilarity | Object | right PHC | left angular | -0.39 | 0.012 |
| SecondSimilarity | Object | right PHC | right supramarginal | -0.39 | 0.014 |
| SecondSimilarity | Object | left PHC | left supramarginal | -0.38 | 0.018 |
| FirstSimilarity | Object | right PHC | right angular | -0.37 | 0.021 |
| SecondSimilarity | Object | left PRC | left supramarginal | -0.36 | 0.023 |
| FirstSimilarity | Object | right PHC | left middle frontal | -0.34 | 0.032 |
| FirstSimilarity | Object | left PHC | right angular | -0.34 | 0.033 |
| SecondSimilarity | Object | left PHC | right supramarginal | -0.33 | 0.037 |
| FirstSimilarity | Object | right PRC | right middle frontal | -0.33 | 0.038 |
| SecondSimilarity | Object | left PHC | left middle frontal | -0.33 | 0.04 |
| FirstSimilarity | Object | left PRC | left angular | -0.32 | 0.044 |
| FirstSimilarity | Object | left PHC | left angular | -0.32 | 0.049 |

**Supplementary Table 13. Location mnemonic discrimination and frontoparietal-medial temporal connectivity**

| Similarity Measure | Task | MTL ROI | Frontoparietal ROI | Spearman r | p |
|---|---|---|---|---|---|
| FirstSimilarity | Location | left PRC | right middle frontal | -0.38 | 0.017 |
| FirstSimilarity | Location | right PHC | right angular | -0.35 | 0.03 |
| FirstSimilarity | Location | right PHC | right middle frontal | -0.32 | 0.045 |

### Supplementary Methods

#### Stimuli assignment to conditions

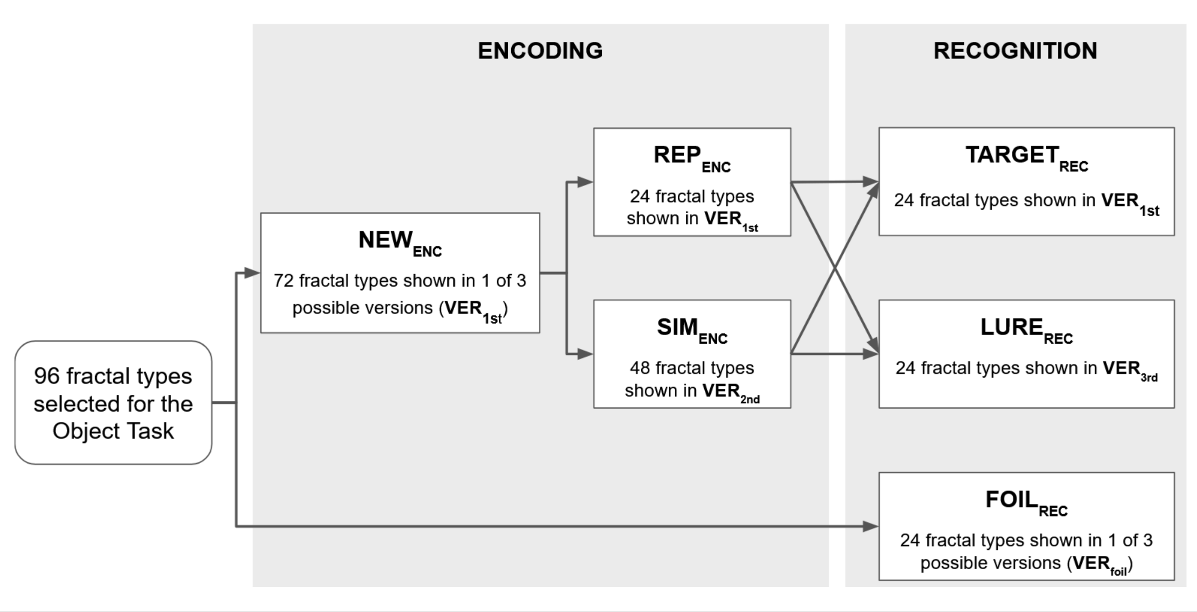

**Supplementary Figure 5. Structure of the Object task**

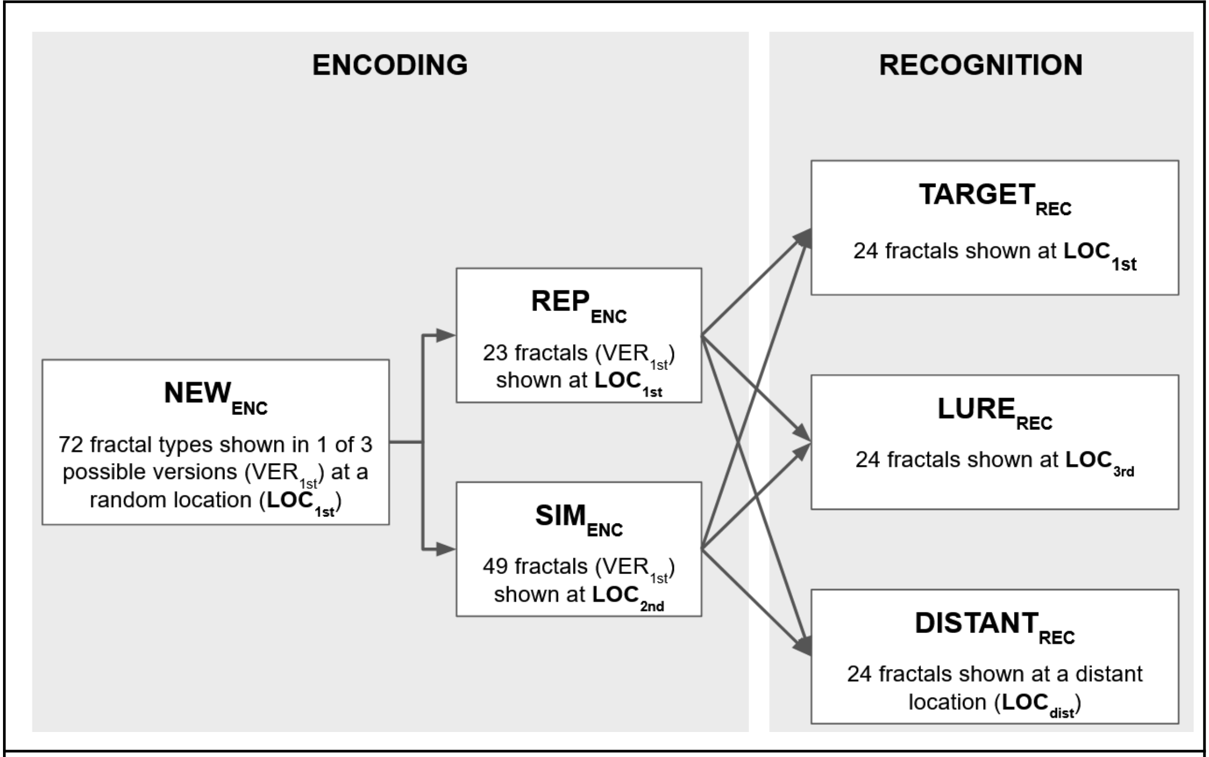

**Supplementary Figure 6. Structure of the Location task**

#### Spatial Similarity Manipulation

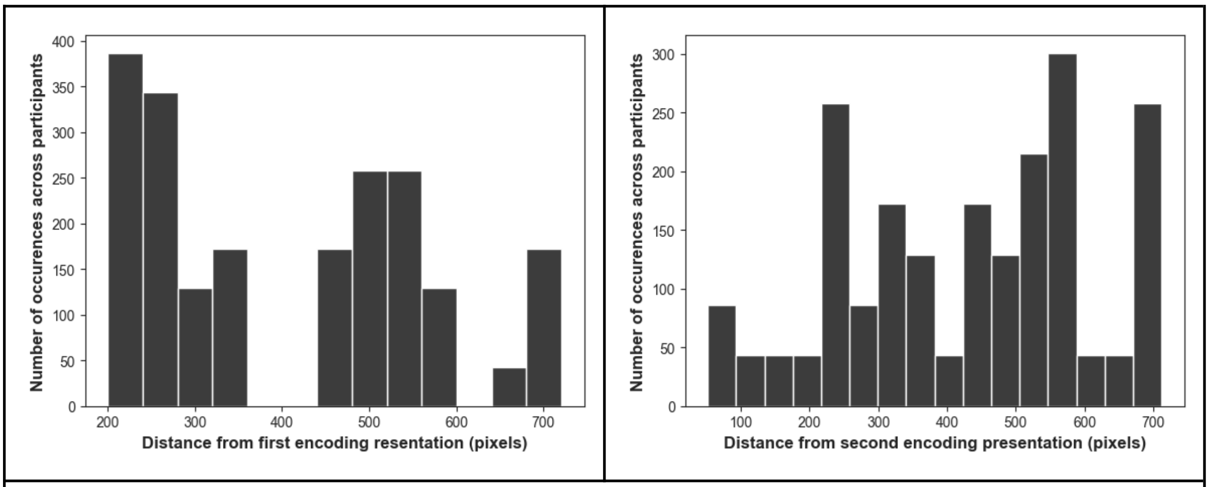

**Supplementary Figure 7.**

#### Object Similarity Rating Experiment

Thirty participants were recruited from the local student community to rate the similarity between the three versions of fractal types. Participants were screened for color blindness with a short Ishihara Test (Ishihara, 1951) consisting of 5 plates. The experiment was programmed with Psytoolkit 3.4.0 (Stoet, 2010, 2017). Participants were randomly assigned to 3 groups, and each group rated a non-overlapping set of fractal pairs, so that overall the three groups rated all pairs of fractal versions within each fractal type (e.g., Group A rated the pair 1a - 1b, Group B rated 1b - 1c, and Group C rated 1a - 1c). In each trial, participants were presented with a pair of images, showing 2 different versions of the same fractal type. Participants had a maximum of 10 seconds to rate the similarity of the two images on a scale from 0 (identical) to 9 (very different). Rating trials were preceded by 20 practice trials, after which participants received feedback on their average rating, and were encouraged to use the whole extent of the scale. Participants completed 200 rating trials. In 19 of the rating trials, 2 identical fractal images were shown, which were not used as stimuli in the mnemonic discrimination task. The experimental session lasted 15-20 minutes, and participants were compensated with course credits.

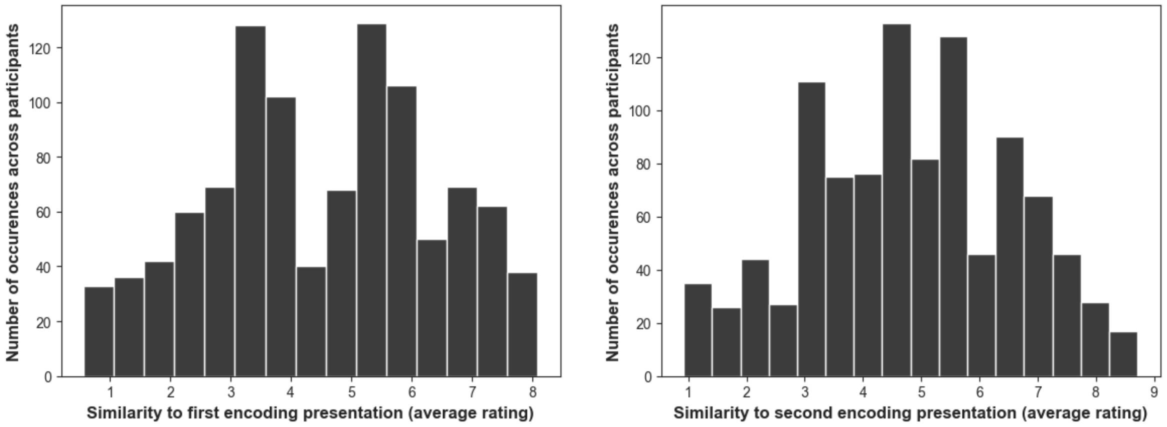

**Supplementary Figure 8.**

#### Segmentation of region of interests in the medial temporal lobe

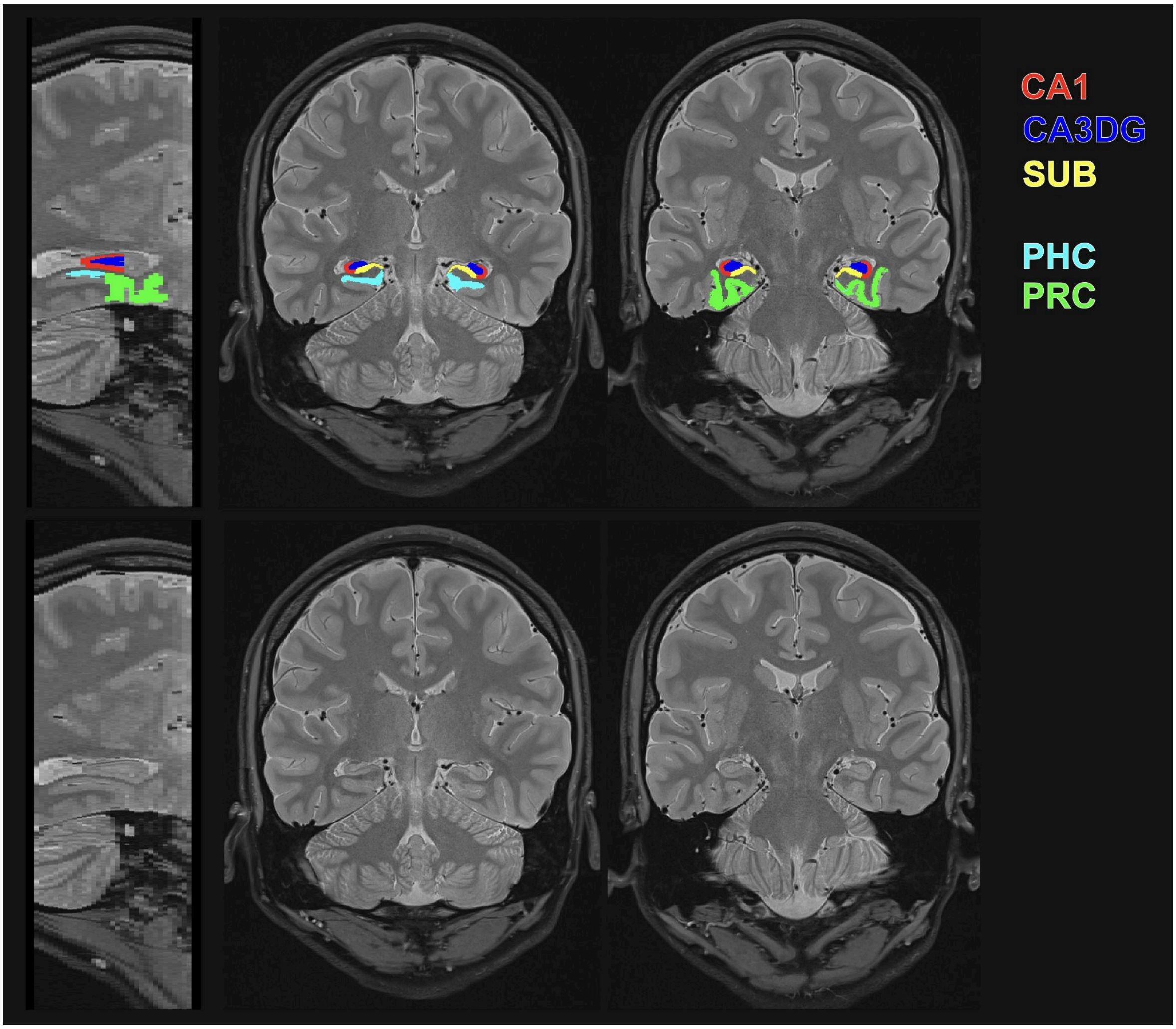

**Supplementary Figure 9. Medial temporal ROIs on an example segmentations**

**Registration of partial FOV high-resolution PD images to functional scans**

We used the following three ANTs calls.

Step 1. Individual T1 weighted images were registered to the partial FOV high resolution PD images with the ANTs call:

```
antsRegistrationSyN.sh \
  -d 3 \
  -f highres_PD.nii.gz \
  -m T1.nii.gz \
  -t a \
  -o T1-To-hPD_
```

Step 2. The mean of the first Object Encoding functional run was registered to the individual T1 weighted anatomy with the ANTs command:

```
antsRegistration \
  --dimensionality 3 \
  --float 0 \
  --output [meanFunc-To-T1_, meanFunc-To-T1_Warped.nii.gz] \
  --interpolation Linear \
  --winsorize-image-intensities [0.005,0.995] \
  --use-histogram-matching 0 \
  --initial-moving-transform [T1.nii.gz, meanFunc.nii.gz, 1] \
  --transform Rigid[0.1] \
  --metric MI[T1.nii.gz, meanFunc.nii.gz,1,32,Regular,0.20] \
  --convergence [1000x500x100x0,1e-6,10] \
  --shrink-factors 8x4x2x1 \
  --smoothing-sigmas 3x2x1x0vox \
  --transform SyN[0.1,3,0] \
  --restrict-deformation 1x1x0 \
  --metric CC[T1.nii.gz, meanFunc.nii.gz,1,4] \
  --convergence [10,1e-6,10] \
  --shrink-factors 1 \
  --smoothing-sigmas 0vox  \
  --x MTL_mask.nii.gz \
```

Step 3. Medial temporal ROI masks were transformed to the functional space using the transformation matrices and warp image resulting from the above registrations:

```
antsApplyTransforms \
  -d 3 \
  -i ROI_mask.nii.gz\
  -t [meanFunc-To-T1_0GenericAffine.mat, 1] \
  -t meanFunc-To-T1_1InverseWarp.nii.gz \
  -t [T1-To-hPD_0GenericAffine.mat, 1] \
  -n GenericLabel \
  -o ROI-To-meanFunc.nii.gz
```

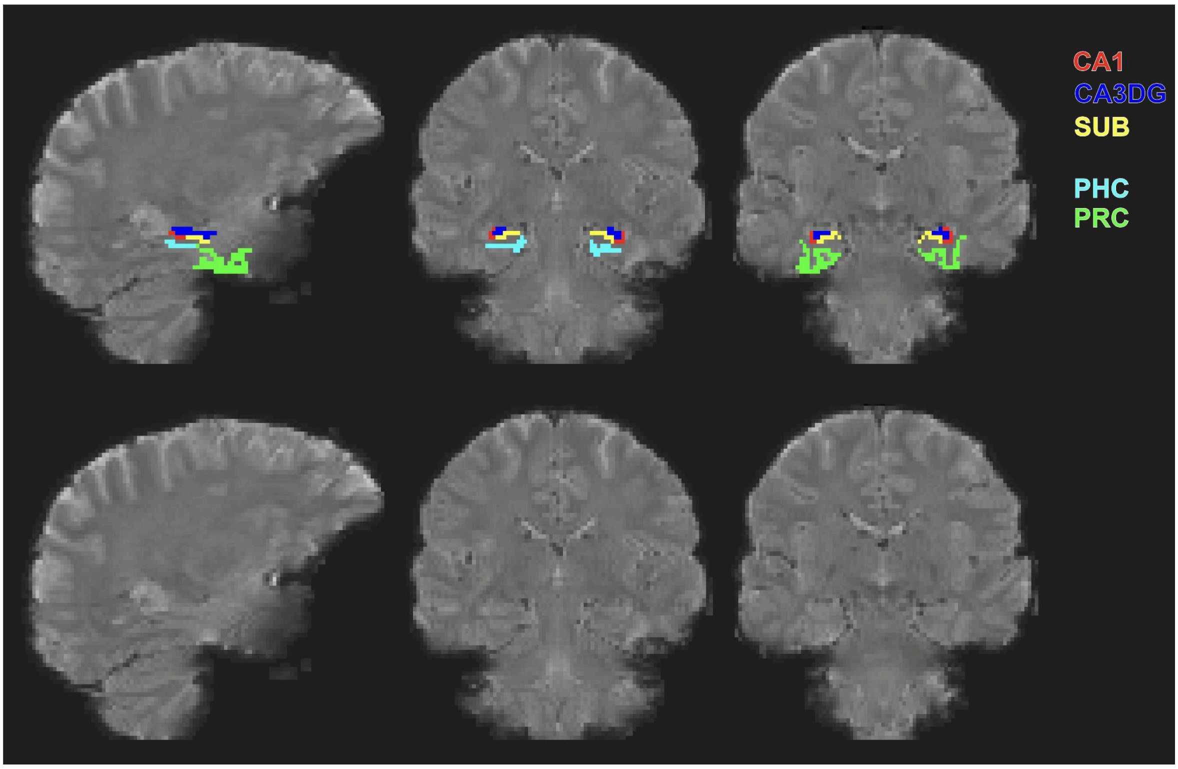

**Supplementary Figure 10. Example ROI to Functional registrations**

### Mixed effects model assumptions

Model assumptions were tested using the performance 0.12.3 (Lüdecke et al., 2021) and EnvStat 3.0.0 (Millard, 2013) packages in R.

#### Behavioral Models: Generalized  binomial mixed effects models

In the behavioral analysis, all modeled responses were binary (new response: 0/1). No outliers were found that have biased the models. Binned residual checks indicated that over 80% of residuals of all models were inside the error bounds, except for Model B of the Object Task (78% of residuals were inside the error bounds). We found no significant overdispersion in any of the models.

#### Neural Models: Linear mixed effects models

Q-Q Plots and residual plots indicated that residuals were normally distributed with homogenous variance across groups (Suppl. Fig 9).

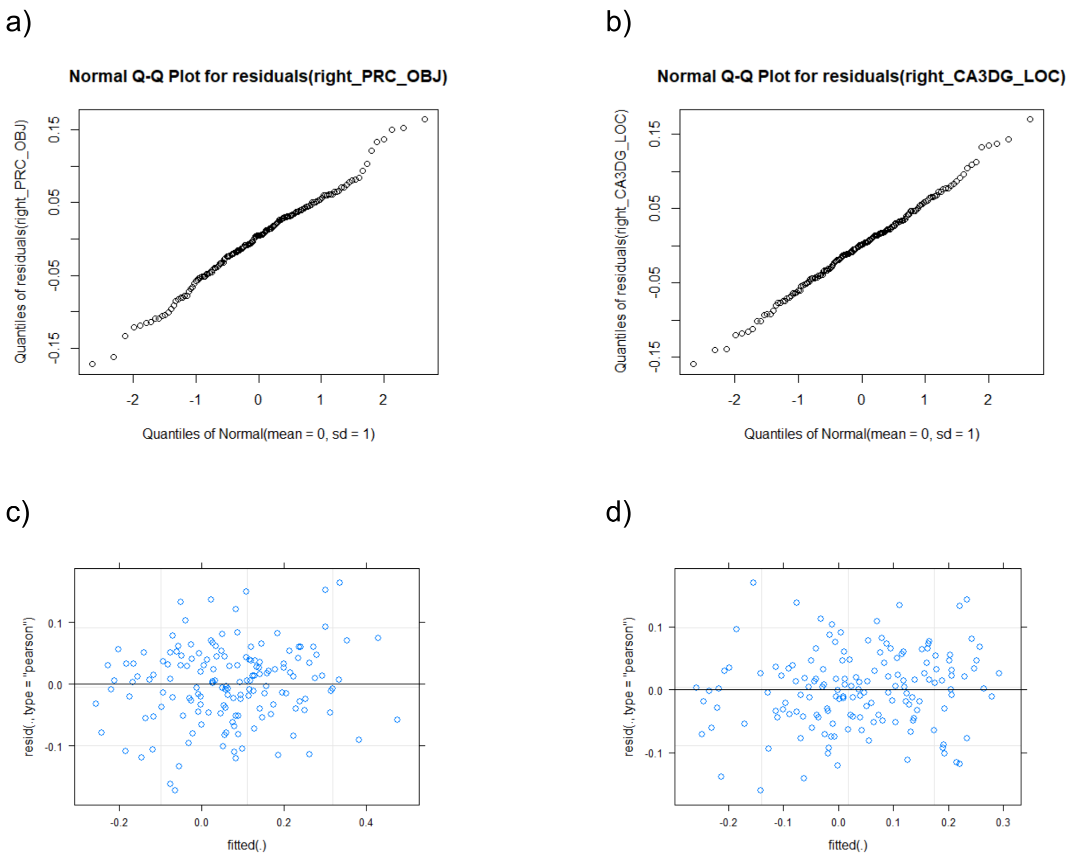

**Supplementary Figure 11.**
Q-Q Plots indicated that the residuals from models relating neural responses to similarity were normally distributed. a) Q-Q plot of the model predicting the response of the right PRC based on object similarity. b) Q-Q plot of the model predicting the response of the right CA3DG based on spatial similarity. Plotting standardized residuals against the predicted values suggests that the assumption  of homoscedasticity was met in both models. c) Residuals from the right PRC response ~ object similarity model. d)  Residuals from the right CA3DG response ~ spatial similarity model.

### References

Ishihara, S. (1951). *Tests for colour-blindness*. Kanehara Tokyo.

Lüdecke, D., Ben-Shachar, M. S., Patil, I., Waggoner, P., & Makowski, D. (2021). performance: An R Package for Assessment, Comparison and Testing of Statistical Models. *Journal of Open Source Software*, *6*(60), 3139. https://doi.org/10.21105/joss.03139

Millard, S. P. (2013). *EnvStats: An R Package for Environmental Statistics*. Springer. https://www.springer.com

Stoet, G. (2010). PsyToolkit: A software package for programming psychological experiments using Linux. *Behavior Research Methods*, *42*, 1096–1104.

Stoet, G. (2017). PsyToolkit: A novel web-based method for running online questionnaires and reaction-time experiments. *Teaching of Psychology*, *44*(1), 24–31.
